# Deconvolving feeding niches and strategies of abyssal holothurians from their stable isotope, amino acid, and fatty acid composition

**DOI:** 10.1101/2023.02.07.527477

**Authors:** Tanja Stratmann, Peter van Breugel, Dick van Oevelen

**Affiliations:** NIOZ Royal Netherlands Institute for Sea Research, Department of Ocean Systems, ‘t Horntje (Texel), The Netherlands; Utrecht University, Department of Earth Sciences, Utrecht, The Netherlands; NIOZ Royal Netherlands Institute for Sea Research, Department of Estuarine and Delta Systems, Yerseke, The Netherlands

**Keywords:** Echinodermata, NLFA, PLFA, sea cucumber, diet, deep sea

## Abstract

Holothurians are the dominant megabenthic deposit feeders in the Peru Basin (South-East Pacific) and feed to various degrees of selectively on the heterogenous pool of sedimentary detritus, but diet preferences for most holothurian species are unknown. This study reconstructs the diets of 13 holothurian species of the orders Elasipodida, Holothuriida, and Synallactida, from bulk stable isotope analyses (δ^13^C, δ^15^N) of holothurian body walls and guts, gut contents, and feces that were combined with compound-specific stable isotope analyses of amino acids, phospholipid-derived fatty acids, and neutral lipid-derived fatty acids in the body wall. Fatty acid concentrations showed high levels of storage lipids, an likely adaption to limited food supply to abyssal plains. Amino acid δ^15^N isotope values allowed estimating trophic levels of holothurian species and calculating heterotrophic re-synthesis of amino acids. Fatty acids served as trophic markers for feeding on diatom- and dinoflagellate derived phytodetritus, bacteria, Foraminifera, and detritus containing the PUFA C22:1ω9-*cis*. Several holothurian species seemed to be secondary consumers of detritus, while bacteria in their guts were primary consumers of this detritus. A Sørensen–Dice coefficient based cluster analysis using data of trophic levels, levels of heterotrophic re-synthesis of amino acids, feeding selectivity, and food sources/ diet suggested three trophic groups, characterized by different trophic levels. We show that this multi-biomarker driven approach allows to deconvolve trophic niches and feeding selectivity in one of the most challenging environments on earth and to identify dependence of deep-sea species to organic matter inputs that vary with season and/or climate.

## Introduction

Holothurians are one of the most abundant epifauna in the deep sea (Billett et al. 2001; Ruhl 2007; Alt et al. 2013; Stratmann et al. 2018) and they can be suspension and deposit feeders (Massin 1982). On soft sediment, deposit feeding holothurians either dig into the sediment as funnel-feeder or conveyor belt-feeder or scavenge the surface sediment as rake feeders (Massin 1982). In this way, they take up particulate organic matter that is deposited on or buried in the sediment (Roberts et al. 2000). Holothurians selectively feed on the organic sources in the sediment. The analysis of gut contents from holothurians collected at the Porcupine Abyssal Plain (PAP, NE Atlantic) showed that e.g. *Amperima rosea* Perrier, 1886, *Peniagone diaphana* Théel, 1882, and *Oneirophanta mutabilis mutabilis* Théel, 1879, feed selectively on fresh phytodetritus (FitzGeorge-Balfour et al. 2010). However, when fresh phytodetritus is scarce, *O. mutabilis mutabilis* feeds on more refractory detritus material (FitzGeorge-Balfour et al. 2010) which is primarily consumed by the microbial community in its gut (Romero-Romero et al. 2021). Other species have a less selective feeding behavior, e.g. *Psychropotes longicauda* Théel, 1882, *Molpadiodemas villosus* Théel, 1886, and *Molpadia blakei* Théel, 1886, (FitzGeorge-Balfour et al. 2010). Though it seems that feeding selectivity and diet preferences of holothurians are well known, this is actually true for very few species. For most abyssal holothurians, in fact, these information are very rudimentary (e.g. Billett 1991; Roberts et al. 2000)).

Whereas holothurians alter the chemical composition of detritus in the sediment, this detritus composition also affects the species composition of holothurians (Wigham et al. 2003; FitzGeorge-Balfour et al. 2010). At PAP, especially *A. rosea, P. diaphana* and *O. mutabilis mutabilis* had a high concentration of carotenoids in their ovaries which are important for the reproductive success of the species (Tsushima 2007; Svensson and Wong 2011). Therefore, (Wigham et al. 2003) suggested that higher concentrations of carotenoids in the gonads of *A. rosea* as compared to other holothurians might give this species a reproductive advantage which could explain the so-called ‘Amperima’ event. During this event, the density of *A. rosea* increased by three orders of magnitude due to large-scale recruitment events that followed changes in the organic carbon flux to the abyssal plain, even though the total megafauna biomass did not change significantly (Billett et al. 2010).

Amino acids, the building stones of proteins, are required to produce enzymes, structural tissue of fauna, and cell walls of bacteria (Phillips 1984; Libes 2009). Half of the 20 most common amino acids in faunal proteins can be synthesized by the organism itself (Phillips 1984), whereas the other half has to be taken up with the diet and are therefore called ‘essential’ amino acids (EAA) (Phillips 1984). Amino acids include ‘source amino acids’ (i.e., glycine, serine, phenylalanine, tyrosine, lysine), which preserve their δ^15^N values along the trophic chain because no new bonds are formed to the N atom nor are bonds cleaved (Chikaraishi et al. 2009). Other amino acids are ‘metabolic amino acids’ (i.e., theorine) and ‘trophic amino acids’ (i.e., asparagine, glutamine, alanine, isoleucine, leucine, valine, proline). The δ^15^N values of ‘trophic amino acids’ become enriched during metabolic transamination when nitrogen bonds are cleaved (Chikaraishi et al. 2009). The larger the difference between the ‘source amino acids’ and the ‘trophic amino acids’, the higher is the trophic level of an organism, so the ratio of the δ^15^N values of glutamic acid and phenylalanine has be used to estimate the trophic level of an organism following (Chikaraishi et al. 2009).

Fatty acids, the main components of lipids, serve as energy source, are involved in the transduction of signals, in gene expression, and are components of membranes (Burdge and Calder 2014). They contain neutral lipid-derived fatty acids (NLFAs) and phospholipid-derived fatty acids (PLFAs) (Dalsgaard et al. 2003). NLFAs are required to build wax esters and the storage lipids triacylglycerols, whereas PLFAs are necessary to build structural phospholipids of cell membranes (Dalsgaard et al. 2003). Fatty acids may be unsaturated or saturated and generally a higher number of unsaturated bonds implies that the fatty acid is more labile than a fatty acid with fewer unsaturated bonds (Pond et al. 1997). ‘Essential’ fatty acids have to be taken up with the diet because they can generally only be synthesized *de novo* by primary producers (Dalsgaard et al. 2003), except for a few hydrothermal vent shrimp species and worms that are also able to synthesize them (Pond et al. 1997, 2002). Since several fatty acids are transferred conservatively (i.e., untransformed) from primary producers and primary consumers to higher trophic levels, they may serve as trophic markers and inform about diets (Dalsgaard et al. 2003).

To decipher feeding types and diet preferences of holothurians from the Peru Basin, compound-specific stable isotope analyses of bulk tissue, gut content, and feces were combined with compound-specific stable isotope analysis of amino acids and fatty acids. We addressed the following research questions: (1) Do the holothurian species have different trophic levels? (2) Can specific feeding strategies and diet preferences identified for the different species?

## Material and methods

### Sampling of holothurians

Holothurians of the putative species Elpidiidae gen sp. Théel, 1882 (n = 1), *Amperima* sp. Pawson, 1965 (n = 4), *Benthodytes* sp. Théel, 1882 (n = 2), *Benthodytes typica* Théel, 1882 (n = 1), *Galatheathuria* sp. Hansen & Madsen, 1956 (n = 1), *Oneirophantha* sp. Théel, 1879 (n = 1), *Psychronaetes hanseni* Pawson, 1983 (n = 1), *P. longicauda* (n = 1), *Psychropotes semperiana* Théel, 1882 (n = 1), *Synallactes* sp. Ludwig, 1894 (morphotype “pink”; n = 1), and Synallactidae gen sp. Ludwig, 1894 (n = 2) were collected opportunistically with the ROV with the ROV suction sampler in the Peru Basin (Table 1). As a result, sampling of several species was not balanced, but due to logistical constraints it was often limited to n = 1 or n = 2. Aboard RV *Sonne*, the length, height, and width of each holothurian specimen was measured and the specimens were dissected to separate the gut and its content from the remaining tissue. All samples were shock-frozen in liquid nitrogen and stored frozen at -20°C.

**Table 1.**
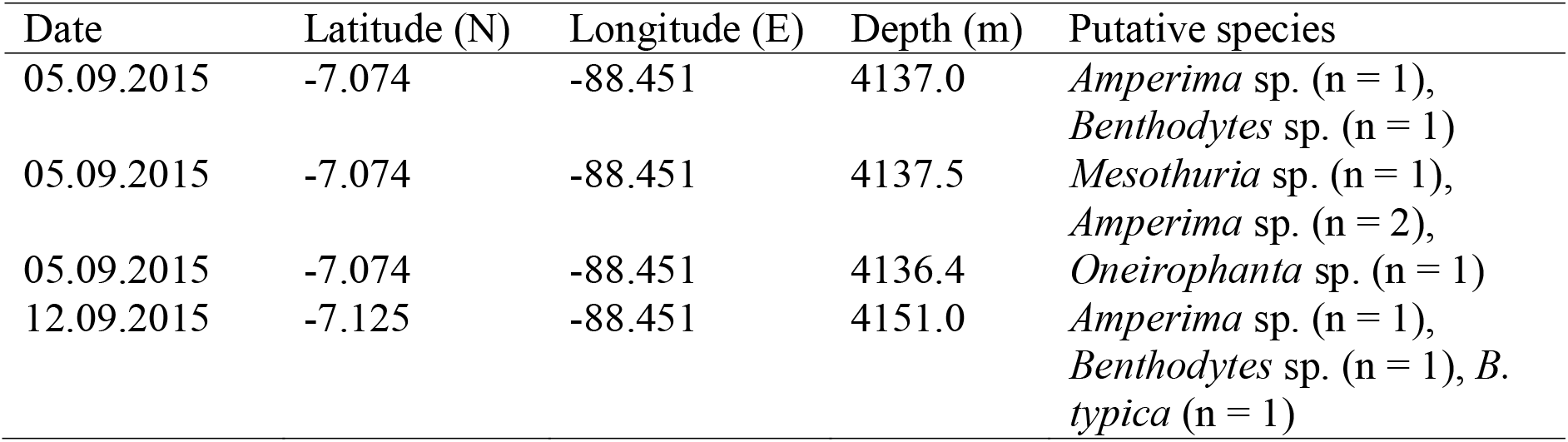
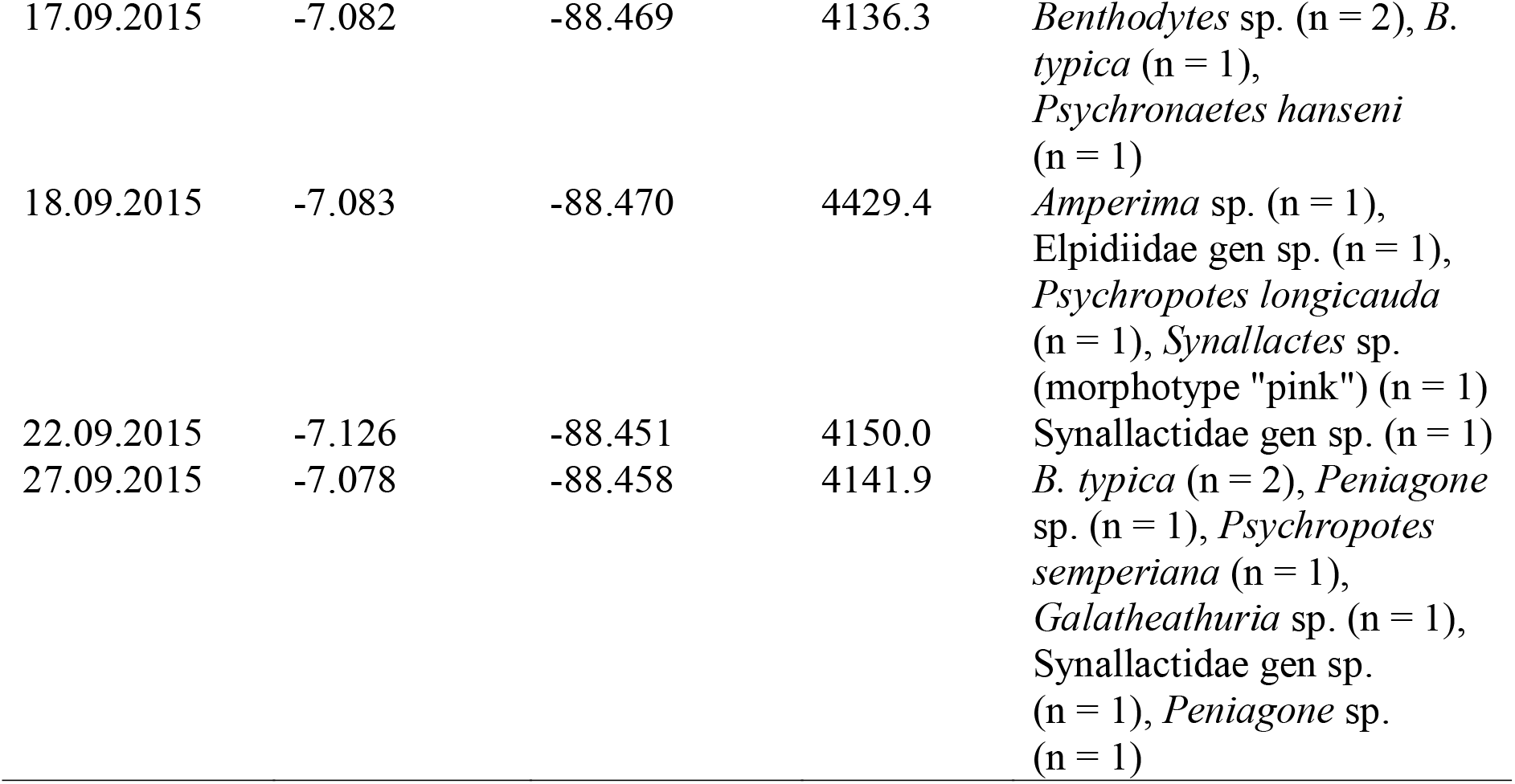
Details of sampling location and collected holothurian specimens from RV *Sonne* research cruise SO242-2.

Additionally, the putative holothurians species *Amperima* sp. (n = 3), *Benthodytes* sp. (n = 3), *B. typica* (n = 4), *Mesothuria* sp. Ludwig, 1894 (n = 1), *Peniagone* sp. Théel, 1882 (n = 1), and Synallactidae gen sp. Ludwig, 1894 (n = 1) from the study of (Brown et al. 2018) were used. These specimens were collected with the ROV suction sampler in the Peru Basin and transported to respiratory chambers to measure oxygen consumption of individual holothurian specimen over a period of 72 hours. Aboard RV *Sonne*, the holothurians specimens were measured (length, height, width), shock-frozen intact in liquid nitrogen, and stored at. Feces of holothurians that defecated inside the respiratory chambers were sampled and frozen at -21°C.

In the shore-based laboratory at NIOZ-EDS (Yerseke, Netherlands), the samples were freeze-dried and finely-ground with mortar and pestle. The organic (org.) C/ δ^13^C and N/ δ^15^N content of the holothurian tissue and of the acidified holothurian gut content were measured with a Thermo Flash EA 1112 elemental analyzer (EA; Thermo Fisher Scientific, USA) which was coupled to a DELTA V Advantage Isotope Ratio Mass Spectrometer (IRMS; Thermo Fisher Scientific, USA). Stable isotope values are presented in δ notation relative to Vienna Pee Dee Belemnite for δ^13^C and relative to air for δ^15^N.

Sediment grain size of holothurian gut content was determined by laser diffraction on freeze-dried and sieved (<1 mm) sediment samples in a Malvern Mastersizer 2000.

### Analysis of amino acids

Total hydrolysable amino acids (THAA) from holothurian tissue were extracted following a modified protocol of Veuger et al. (2005): Briefly, THAAs in holothurian tissue were hydrolyzed by adding 0.01 to 0.02 g freeze-dried finely ground tissue to 1.5 ml 6 M HCl in 10 ml screw-cap vials. A N_2_-headspace was created in the vials by flushing with N_2_-gas for 10 sec before the vials were closed and heated for 20 h at 110°C. After cooling, 10 μL internal L-Norleucine standard per mg dry faunal tissue (stock solution: 2.5 mg mL^-1^ L-Norleucin acidified with 100 μL 12 M HCl) was added and the solution was evaporated under N_2_-flow at 60°C. THAAs from holothurian tissue were derivatized by adding 0.5 ml acidified propan-2-ol to the sample and by heating the closed vials at 110°C for 90 min. Afterwards, the vials were cooled down and the solution was evaporated under N_2_-flow at 50°C. After evaporating all solution, 200 μL dichloromethane (DCM) was added and the solution was evaporated again. When the samples were dry, 150 μL DCM and 50 μL pentafluoropropionic anhydride were added, the vials were closed and heated for 10 min at 110°C. The solvent was extracted by adding 0.5 mL chlorophorm and 1 ml phosporus-buffer to the sample, shaking it until the lower chloroform fraction was clear and centrifuging the vials with 2,000 rpm for 10 min. The chloroform fraction was transferred to GC vials and evaporated again. When the sample was completely dry, it was dissolved in ethyl acetate. Concentrations (μg C g^-1^ dry mass DM holothurian tissue) and δ^13^C (‰), and δ^15^N (‰) of THAAs were measured with a HP 6890 gas chromatograph (Hewlet Packard/ Agilent, USA) coupled with a DELTA-Plus Isotope Ratio Mass Spectrometer (Thermo Fisher Scientific, USA) on a polar analytical column (ZB5-5MS; 60m length, 0.32mm diameter, 0.25μm film thickness; Phenomenex, USA).

A list with common abbreviation of amino acids and their full name is presented in Table 2.

**Table 2.**
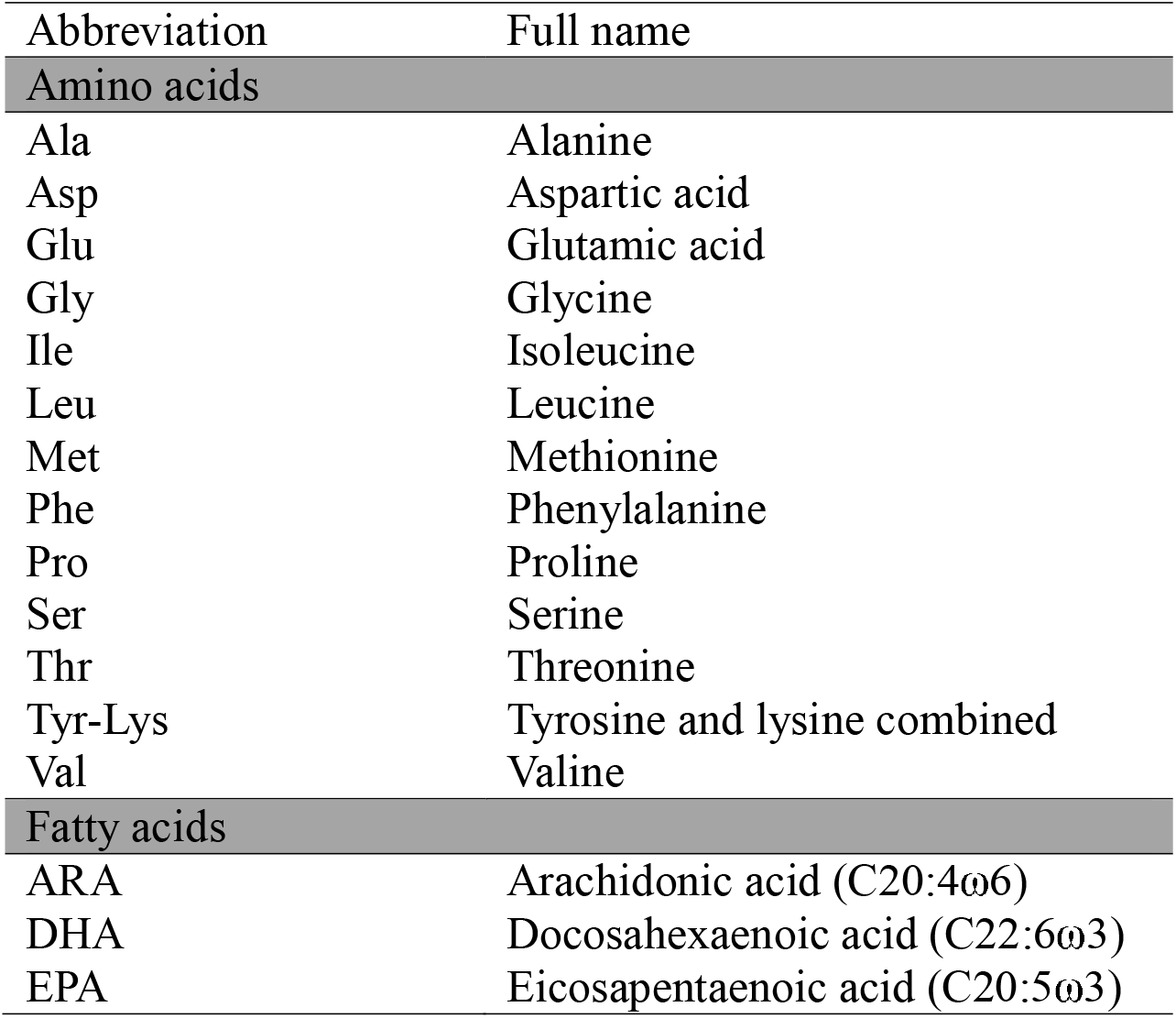
Names and abbreviations of amino acids and fatty acids (PLFAs, NLFAs).

| Abbreviation | Full name |
| --- | --- |
| **Amino acids** |  |
| Ala | Alanine |
| Asp | Aspartic acid |
| Glu | Glutamic acid |
| Gly | Glycine |
| Ile | Isoleucine |
| Leu | Leucine |
| Met | Methionine |
| Phe | Phenylalanine |
| Pro | Proline |
| Ser | Serine |
| Thr | Threonine |
| Tyr-Lys | Tyrosine and lysine combined |
| Val | Valine |
| **Fatty acids** |  |
| ARA | Arachidonic acid (C20:4ω6) |
| DHA | Docosahexaenoic acid (C22:6ω3) |
| EPA | Eicosapentaenoic acid (C20:5ω3) |

### Analysis of fatty acids

Fatty acids (i.e., PLFAs, NLFAs) were extracted from holothurian tissue, feces and gut content following a modified Bligh and Dyer extraction method (Bligh and Dyer 1959; Boschker 2008). Freeze-dried, homogenized powder of holothurian tissue (∼50 – 150 mg), feces and gut content (∼150 mg – 2.0 g) were mixed with 6 ml MilliQ-water 15 ml methanol (HPLC grade, 99.8%), and 7.5% chloroform (HPLC grade, 99.5%) in pre-cleaned test tubes. The tubes were shaken for 2 h, before 7.5 ml chloroform were added and the tubes were shaken again. 7.5 ml MilliQ-water were added and the tubes were stored at -21°C for 12 h for separation of the solvent layers. The lower solvent layer contained the fatty acids extract dissolved in chloroform and was transferred to pre-weighted test tubes. After determining the weight of the chloroform extract, it was fractionated into the different fatty acid classes over an activated silicic acid column (heated at 120ºC for 2 h; Merck Kieselgel 60) via eluting with 7 ml chloroform, 7 ml acetone, and 15 ml methanol. The aceton fraction was discarded, whereas the chloroform fraction containing the NLFAs and the methanol fraction with the PLFAs were collected in separate test tubes and evaporated to dryness.

PLFAs and NLFAs were derivatized to fatty acid methyl esters (FAMEs) by adding 1 ml methanol-toluene mix (1:1 volume/ volume), 20 μl of an internal standard (1 mg 19:0 FAME mL^-1^), and 1 ml 0.2 M metanolic NaOH to the test tubes with the PLFAs and NLFAs extracts. After an incubation at 37ºC for 15 min, 2 ml *n*-hexane, 0.3 ml 1 M acetic acid, and 2 ml MilliQ-water were added. The solution was mixed very well and when the layers had separated, the (top) *n*-hexane layer was transferred to new test tubes. Additional 2 ml *n*-hexane were added to the previously used test tubes that contained the acetic acid-MilliQ-water solution, and the step was repeated. The *n*-hexane layer was transferred again to the new test tubes and 20 μl of a second internal standard (1 mg 12:0 FAME mL^-1^) were added. *n*-hexane was evaporated completely and the FAMEs dissolved in 200 μl *n*-hexane were transferred to measuring vials.

The FAMEs from holothurian tissues were separated on a BPX70 column (50 m length, 0.32 mm inner diameter, 0.25 μm film thickness; SGE Analytical Science) with a HP 6890 gas chromatograph (GC; Hewlet Packard/ Agilent, USA). The FAMEs from feces and gut content were separated on a ZB5-5MS column (60 m length, 0.32 mm diameter, 0.25 μm film thickness; Phenomenex, USA) on the same GC. Concentrations (μg C g^-1^ DM holothurian tissue) and δ^13^C values (‰) of FAMEs in holothurian tissue, feces, and gut content were measured on a Finnigan Delta Plus isotope ratio mass spectrometer (IRMS; Thermo Fisher Scientific, USA) coupled to the GC via a combustion GC-c-III interface (Thermo Fisher Scientific, USA). Identification of peaks of the FAME chromatogram were based on equivalent chain length (ECL) and peak areas were calculated using the two internal standards (12:0 and 19:0) for area correction.

A list with abbreviations and full names of several important fatty acids is presented in Table 2 and Table 3 contains dominant biomarkers.

**Table 3.**
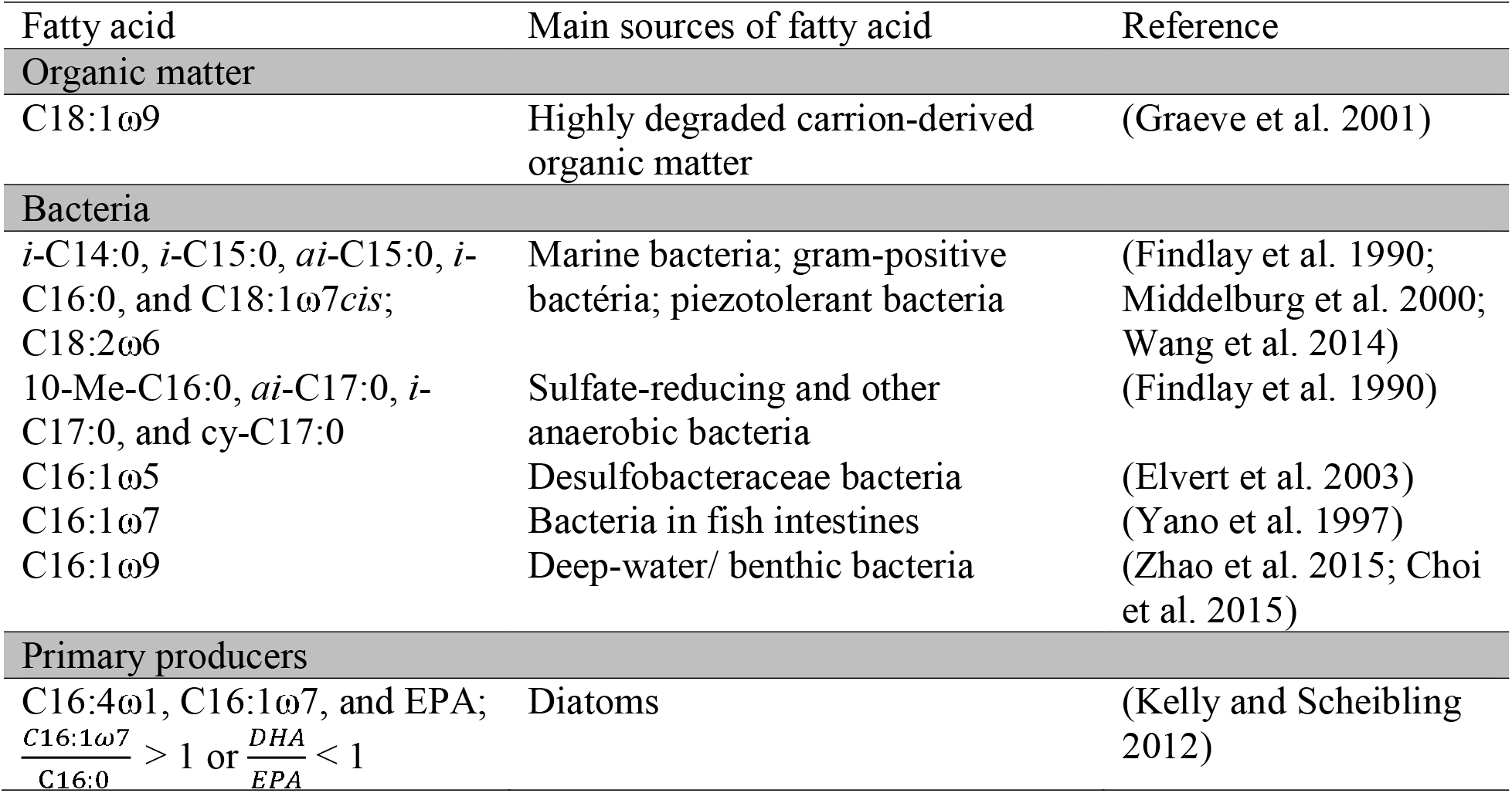
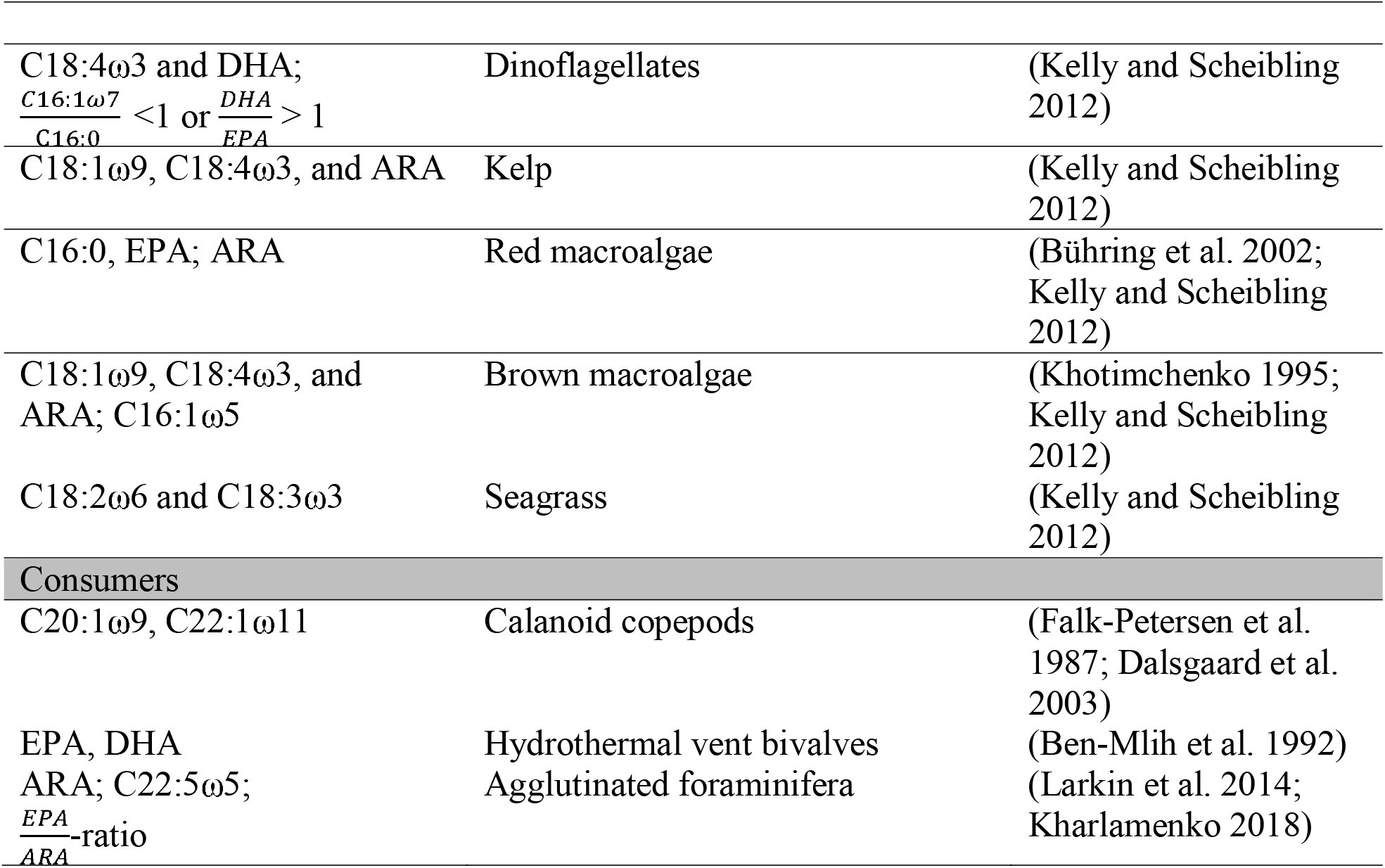
Fatty acids used as biomarkers of potential food sources of holothurians from the Peru Basin.

### Calculations Concentration factors

To examine the degree to which PLFAs were concentrated between surface sediment (0 – 2cm layer; 2.32±0.51 μg C-PLFA g^-1^ DM sediment; Stratmann, unpublished) and gut content and feces, a concentration factor *CF* was calculated:

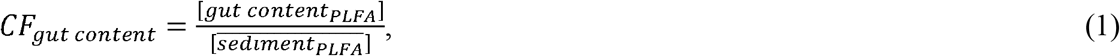

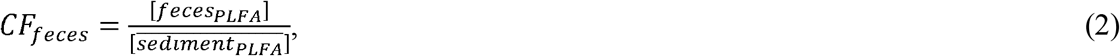

where [gut content_PLFA_] corresponds to the total PLFA concentration in gut content, [feces_PLFA_] to the total PLFA concentration in feces, and 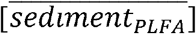 to the average total PLFA concentration in surface sediment.

### Trophic levels

Trophic levels (*TL*) of holothurian species were calculated following (Chikaraishi et al. 2009) as

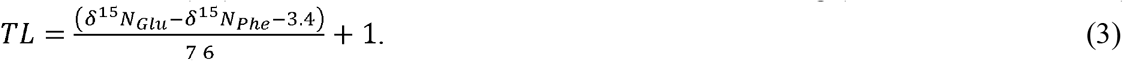

δ^15^*N*_*Glu*_ is the δ^15^N of the amino acid glutamic acid (‰) and δ^*15*^*N*_*Phe*_ corresponds to the δ^15^N of the amino acid phenylalanine (‰).

Trophic levels of two different holothurian species are considered robust, when the difference in trophic levels between two species is >±0.44. This value corresponds to the average standard deviation of the calculated trophic level (σ_*TL*_) across all holothurians that was determined following equation S4 in (Jarman et al. 2017) as:

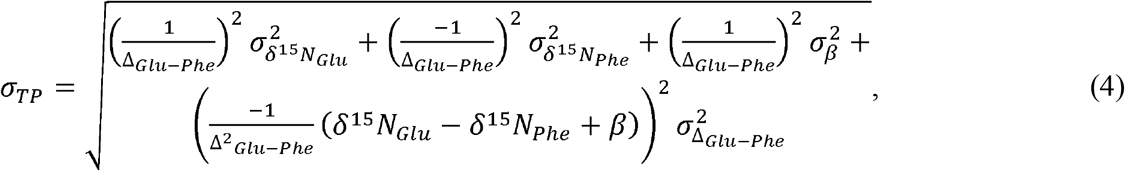

where *σ*_*β*_ is 0.9‰ and *σ*_*Δ*_ is 1.1‰ (Jarman et al. 2017).

### Heterotrophic re-synthesis of amino acids

Total heterotrophic re-synthesis of amino acids (∑*V*) was approximated as the sum of variance of individual δ^15^N values of the trophic amino acids alanine, aspartic acid, glutamic acid, leucine, and proline (McCarthy et al. 2007):

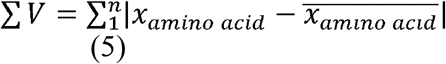

*x* symbolized each trophic amino acid’s δ^15^N value, 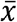 is the average trophic amino acid’s δ^15^N value and *n* is the total number of trophic amino acids used in this calculation (McCarthy et al. 2007).

### Statistics

The Sørensen–Dice coefficient *β*_*sor*_ (Dice 1945; Sørensen 1948; Koleff et al. 2003) was calculated using the ‘betadiver’ function in the *R* (version 4.1.2; R-Core Team 2017) package *vegan* (version 2.6-2; (Oksanen et al. 2017)) to compare holothurian species based on their trophic levels (*TL*), levels of heterotrophic re-synthesis of amino acids (∑*V*), feeding selectivity based on concentration factor (*CF*), and food sources/ diet. For this purpose, the quantitative data *TL* and ∑*V* were first converted into categories (Table 4) and then converted into binary (presence/ absence) data; the categorical data ‘feeding selectivity’ (Table 4) and ‘food/sources diet’ were also converted into binary data. Subsequently, *β*_*sor*_ was clustered by average linkage clustering (unweighted pair-group method using arithmetic averages, UPGMA; Romesburg 1984) using the ‘hclust’ function in *R*. The dendrogram was prepared with R package *factoextra* (version 1.0.7; Kassambara and Mundt 2020).

**Table 4.**
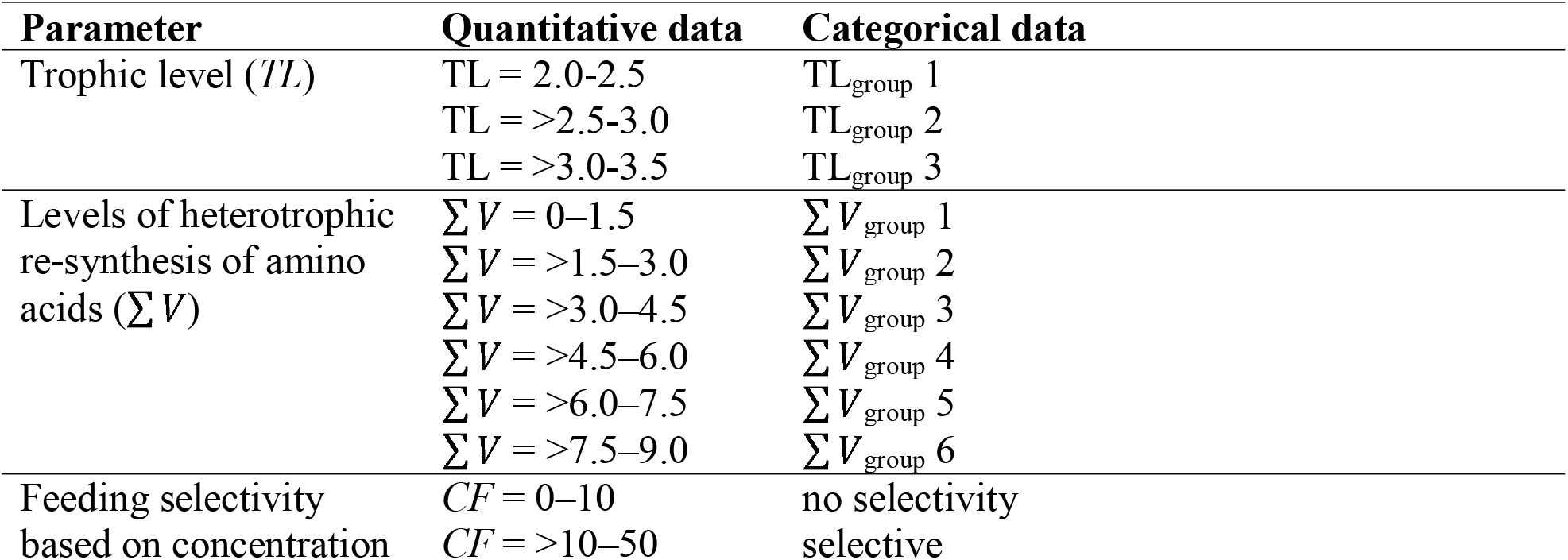
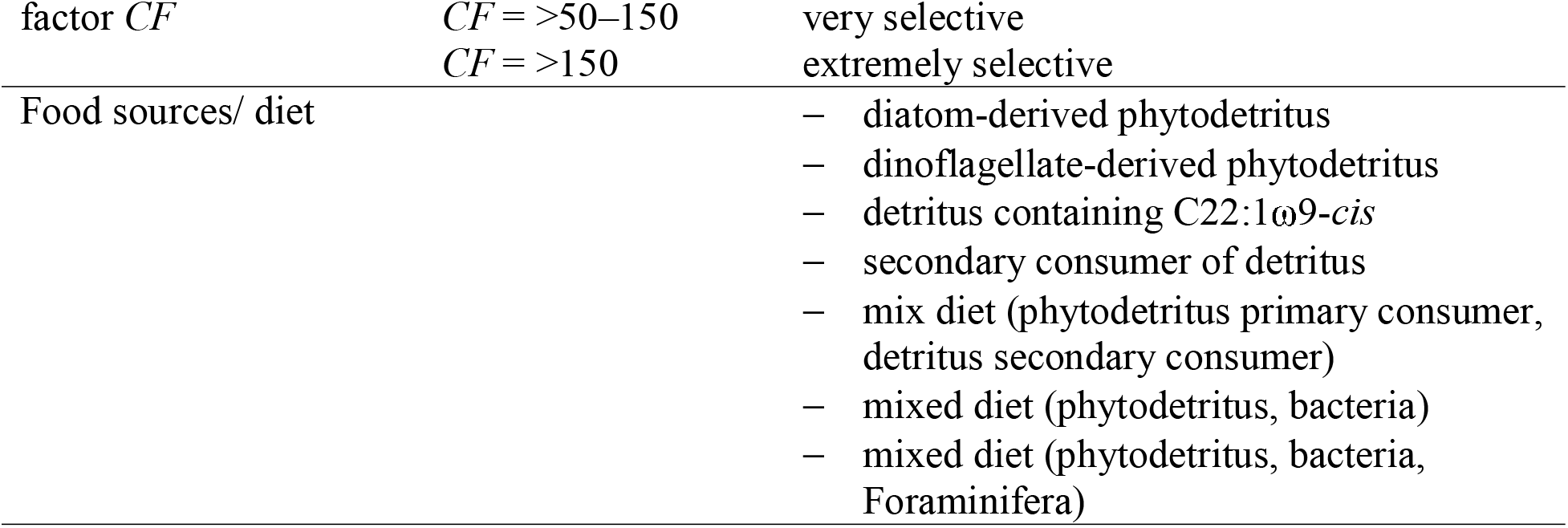
Parameters used to calculate the Sørensen–Dice coefficient *β*_*sor*_ presented as quantitative data (ranges) and categorical data. ‘Food sources/ diet’ includes a list of the main food sources of the investigated holothurian species which were identified by amino acid and fatty acid analysis.

| Parameter | Quantitative data | Categorical data |
| --- | --- | --- |
| Trophic level (*TL*) | *TL* = 2.0-2.5 | *TL*<sub>group 1</sub> |
|  | *TL* = >2.5-3.0 | *TL*<sub>group 2</sub> |
|  | *TL* = >3.0-3.5 | *TL*<sub>group 3</sub> |
| Levels of heterotrophic re-synthesis of amino acids ($\sum V$) | $\sum V$ = 0–1.5 | $\sum V$<sub>group 1</sub> |
| | $\sum V$ = >1.5–3.0 | $\sum V$<sub>group 2</sub> |
| | $\sum V$ = >3.0–4.5 | $\sum V$<sub>group 3</sub> |
| | $\sum V$ = >4.5–6.0 | $\sum V$<sub>group 4</sub> |
| | $\sum V$ = >6.0–7.5 | $\sum V$<sub>group 5</sub> |
| | $\sum V$ = >7.5–9.0 | $\sum V$<sub>group 6</sub> |
| Feeding selectivity based on concentration | *CF* = 0–10 | no selectivity |
|  | *CF* = >10–50 | selective |

| factor *CF* | *CF* = >50–150<br>*CF* = >150 | very selective<br>extremely selective |
| --- | --- | --- |
| Food sources/ diet |  | – diatom-derived phytodetritus<br>– dinoflagellate-derived phytodetritus<br>– detritus containing C22:1ω9-*cis*<br>– secondary consumer of detritus<br>– mix diet (phytodetritus primary consumer, detritus secondary consumer)<br>– mixed diet (phytodetritus, bacteria)<br>– mixed diet (phytodetritus, bacteria, Foraminifera) |

## Results

### Gut content and feces of holothurians

Gut contents of holothurians in the Peru Basin weighed 1.93±3.56 g dry sediment (n = 17) and ranged from 0.11 g dry sediment for *Peniagone* sp. (n = 1) to 12.5 g dry sediment for *P. hanseni* (n = 1; Table 5). Org. C and TN content of the gut content was 5.34±4.13% (n = 17) and 1.04±0.87% (n = 17), respectively, and it contained 244±304 μg C-PLFA g^-1^ DM gut content (n = 8) and 83.3±124 μg C-NLFA g^-1^ DM gut content (n = 10) (Fig. 1a). The concentration factor *CF* for PLFAs in holothurian gut content was on average 105±131 (n = 8) and ranged from 1.17 to 335 for *Benthodytes* sp. (n = 1) and Synallactidae gen sp. (n = 1), respectively (Table 5). The average EPA/ARA-ratio for gut content was 3.20±5.58 (n = 9), the average DHA/EPA-ratio was 0.62±0.68 (n = 9), and the average C16:1ω7/C16:0-ratio was 0.81±0.85 (n = 9) (Fig. 2a).

**Table 5.**
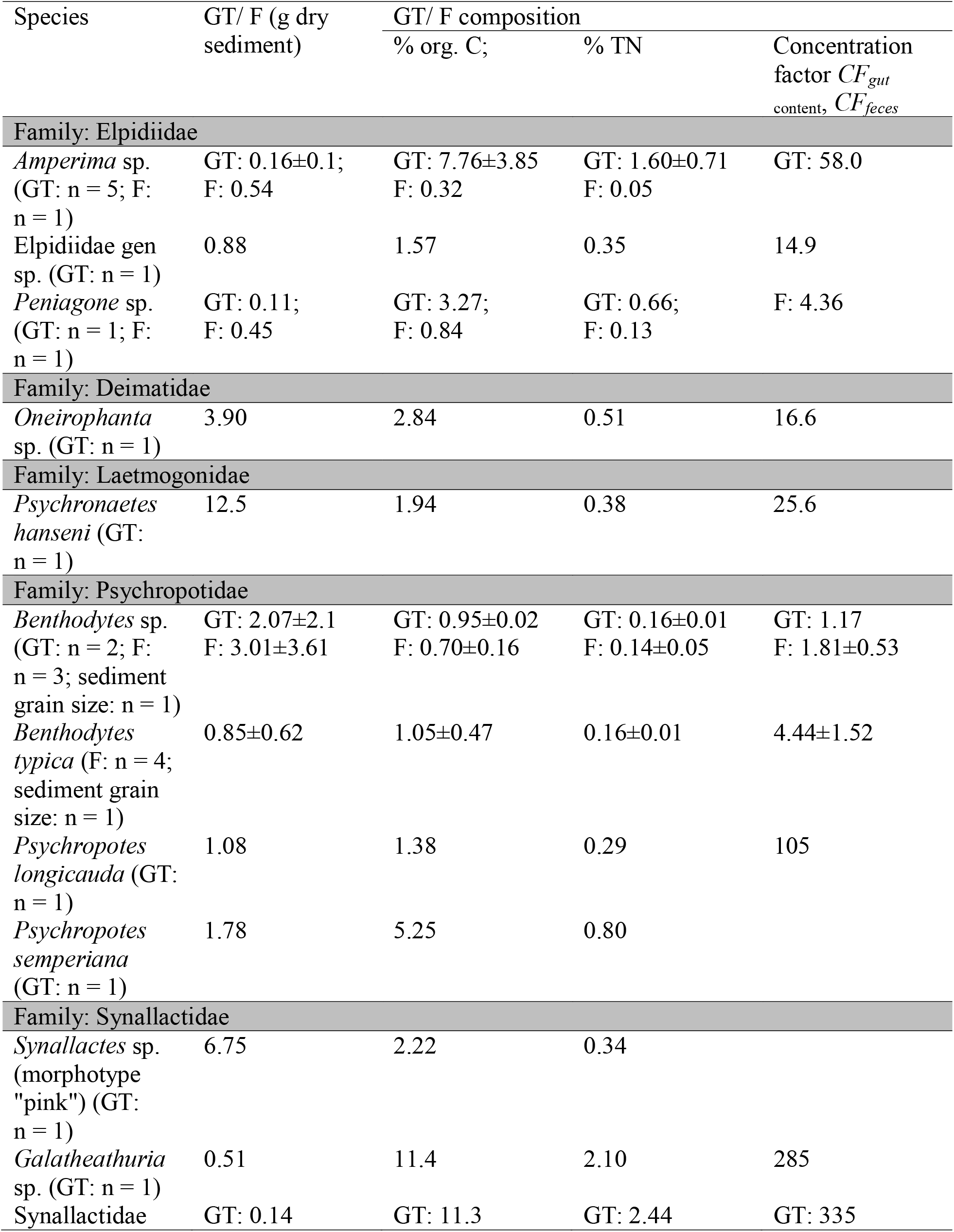
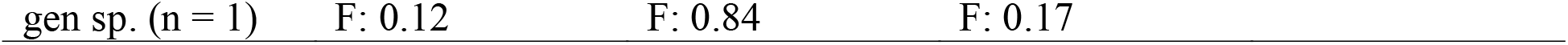
Sedimentological characteristics of holothurian gut content (GT) and feces (F). Data are presented as mean±SD.

**Figure 1.**
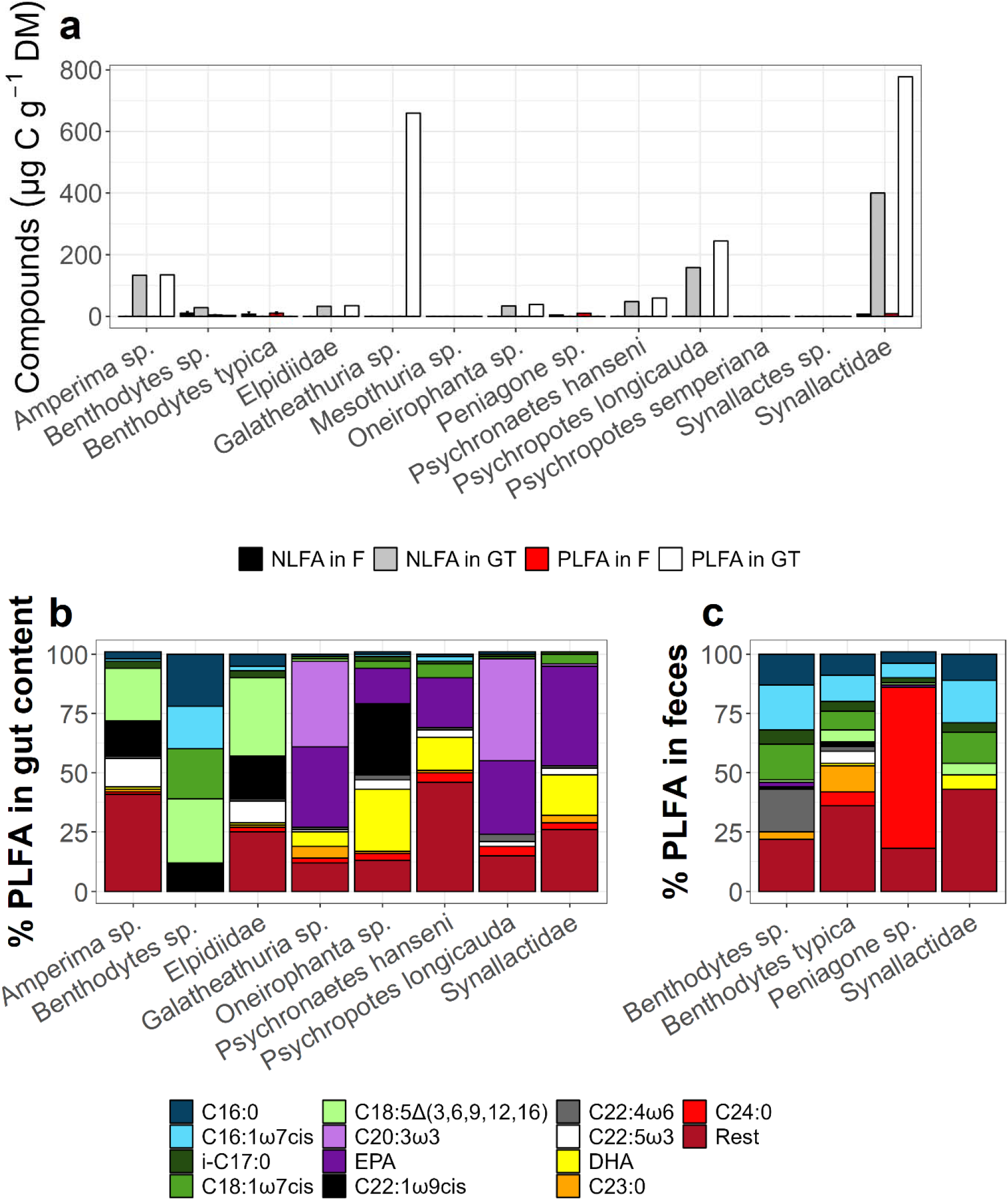
(a) Concentrations (μg C g^-1^ DM. C) of the compounds PLFA and NLFA in holothurian gut content (GT) and feces (F) and the contribution (%) of individual (b, c) PLFAs and (d, e) NLFAs to the total concentrations. The PLFA and NLFA pools ‘Rest’ include all PLFAs and NLFAs, respectively, that contribute <2.5% to total % PLFA and NLFA of the average holothurian gut content/ feces. Error bars in (a) indicate SD.

**Figure 2.**
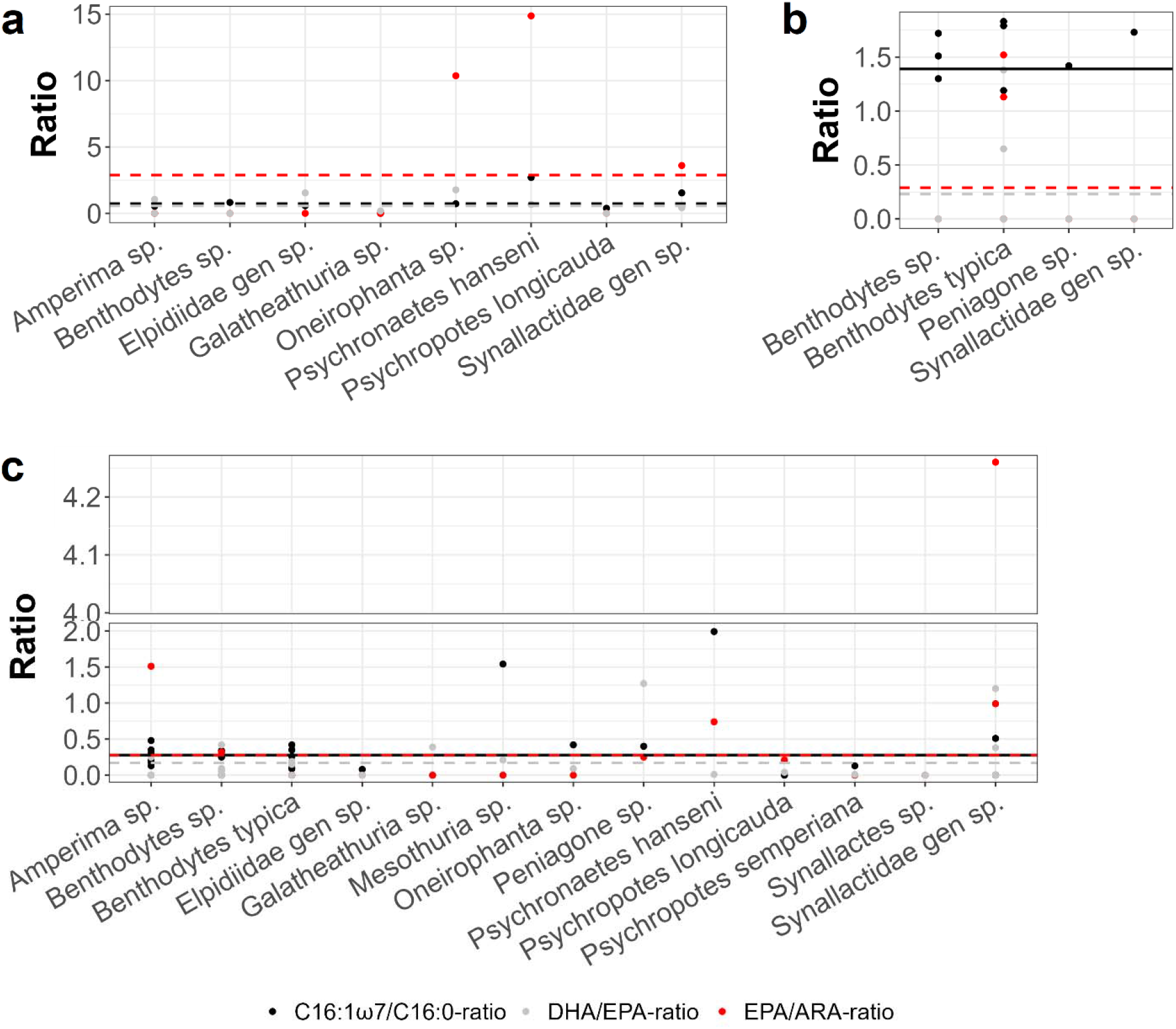
Ratios of C16:1ω7/C16:0, DHA/EPA, and EPA/ARA in (a) holothurian gut content, (b) holothurian feces, and (c) dried holothurian body walls. Horizontal lines show the average value of a ratio based on all samples.

Feces of holothurians weighed 1.36±2.09 g dry sediment (n = 10) and ranged from 0.12 g dry sediment for Synallactidae gen sp. (n = 1) to 3.01±3.61 g dry sediment for *Benthodytes* sp. (n = 3; Table 5). Org. C and TN content of the feces was 0.83±0.37% (n = 10) and 0.14±0.04% (n = 10), respectively, and it contained 7.73±3.60 μg C-PLFA g^-1^ DM sediment (n = 8) and 7.63±5.28 μg C-NLFA g^-1^ DM sediment (n = 15) (Fig. 1a). In holothurian feces (n = 8), PLFAs were on average still 3.33±1.55 times more concentrated compared to the upper 2 cm of sediment (*CF*_*PLFA*_ range: 1.81±0.53 (n = 3) for *Benthodytes* sp. to 4.44±1.52 for *B. typica* (n = 3)) (Table 5). The average EPA/ARA-ratio for feces was 0.29±0.59 (n = 9), the average DHA/EPA-ratio was 0.23±0.48 (n = 9), and the average C16:1ω7/C16:0-ratio was 1.39±0.57 (n = 9) (Fig. 2b).

Gut content and feces consisted to 81.2±%3.73% of silt (grain size: <63 μm; n = 6) and to 1.10% of very fine sand (grain size: 62.5 – 125 μm) (Table 6). The median grain was 15.5±2.27 μm (n = 6).

**Table 6.** Grain size characteristics of gut content (GT) and feces (FC).

| Species | % silt<br>fraction<br>($<63 \mu\text{m}$) | % very<br>fine sand<br>fraction<br>($62.5 - 125 \mu\text{m}$) | % fine<br>sand<br>fraction<br>($125 - 250 \mu\text{m}$) | %<br>medium<br>sand<br>fraction<br>($250 - 500 \mu\text{m}$) | % coarse<br>sand<br>fraction<br>($500 - 1000 \mu\text{m}$) | Median<br>grain<br>size ($\mu\text{m}$) |
| --- | --- | --- | --- | --- | --- | --- |
| **Family: Elpidiidae** |  |  |  |  |  |  |
| Elpidiidae gen<br>sp. (GT; $n = 1$) | 82.3 | 10.0 | 4.91 | 2.05 | 0.92 | 14.7 |
| **Family: Deimatidae** |  |  |  |  |  |  |
| *Oneirophanta*<br>sp. (GT; $n = 1$) | 75.5 | 12.2 | 7.86 | 4.01 | 0.65 | 19.0 |
| **Family: Laetmogonidae** |  |  |  |  |  |  |
| *Psychronaetes*<br>*hanseni* (GT;<br>$n = 1$) | 79.0 | 11.7 | 6.67 | 2.48 | 0.33 | 17.5 |
| **Family: Psychropotidae** |  |  |  |  |  |  |
| *Benthodytes* sp.<br>(F; $n = 1$) | 86.4 | 9.20 | 4.05 | 0.44 | 0.00 | 13.0 |
| *Benthodytes*<br>*typica* (F; $n = 1$) | 81.0 | 10.8 | 6.08 | 1.77 | 0.55 | 14.2 |
| **Family: Synallactidae** |  |  |  |  |  |  |
| Synallactidae<br>gen sp. (F; $n = 1$) | 83.0 | 10.4 | 4.47 | 1.58 | 0.76 | 14.5 |

About 22.3±15.5% (n = 8) of the PLFAs (Fig. 4b) and 23.1±18.4% (n = 7) of the NLFAs (Fig. 4d) found in holothurian gut content consisted of ‘Rest’, i.e., the sum of PLFAs and NLFAs that each contributed <2.5% to total PLFA and NLFA concentrations. The remaining PLFAs consisted to 7.00±6.55% of saturated fatty acids (SFA), 17.0±17.7% monosaturated fatty acids (MUFAs, i.e., fatty acids with one double bond), 10.0±18.4% polyunsaturated fatty acids (PUFAs, i.e., fatty acids with ≥2 double bonds), 41.5±10.6% highly unsaturated fatty acids (HUFAs, i.e., fatty acids with 4 double bonds), and 2.25±1.36% long-chain fatty acids (LCFAs, i.e., fatty acids with ≥24 C atoms). NLFAs included furthermore 36.9±11.7% SFAs, 13.0±11.0% MUFAs, and 27.0±17.3% HUFAs. Feces of holothurian consisted to 29.6±13.3% (n = 8) of the PLFAs category ‘Rest’ (Fig. 4c) and to 31.4±15.6% (n = 8) of the NLFAs category ‘Rest’ (Fig. 4e). The other PLFAs consisted to 20.1±9.58% of SFAs, 25.6±11.7% MUFAs, 14.1±12.4% HUFAs, and 10.6±23.7% LCFAs. The NLFAs included additionally 39.5±11.1% SFAs, 10.3±10.5% MUFAs, and 18.8±24.3% HUFAs.

### Chemical composition of holothurians

Holothurians in the Peru Basin consisted for 93.0±10.2% of water (n = 13) and their dried body walls contained 5.87±3.50% org. C and 1.35±0.80% total N (n = 31), whereas their dried gut tissues consisted of 16.7±8.60% org. C and 3.76±2.18% total N (n = 15). The body wall and gut tissue of the holothurian families Deimatidae and Laetmogonidae had the highest org. C and TN contents, whereas the families Elpidiidae and Psychropotidae had the lowest org. C and TN content in body wall tissue (Table 7).

**Table 7.**
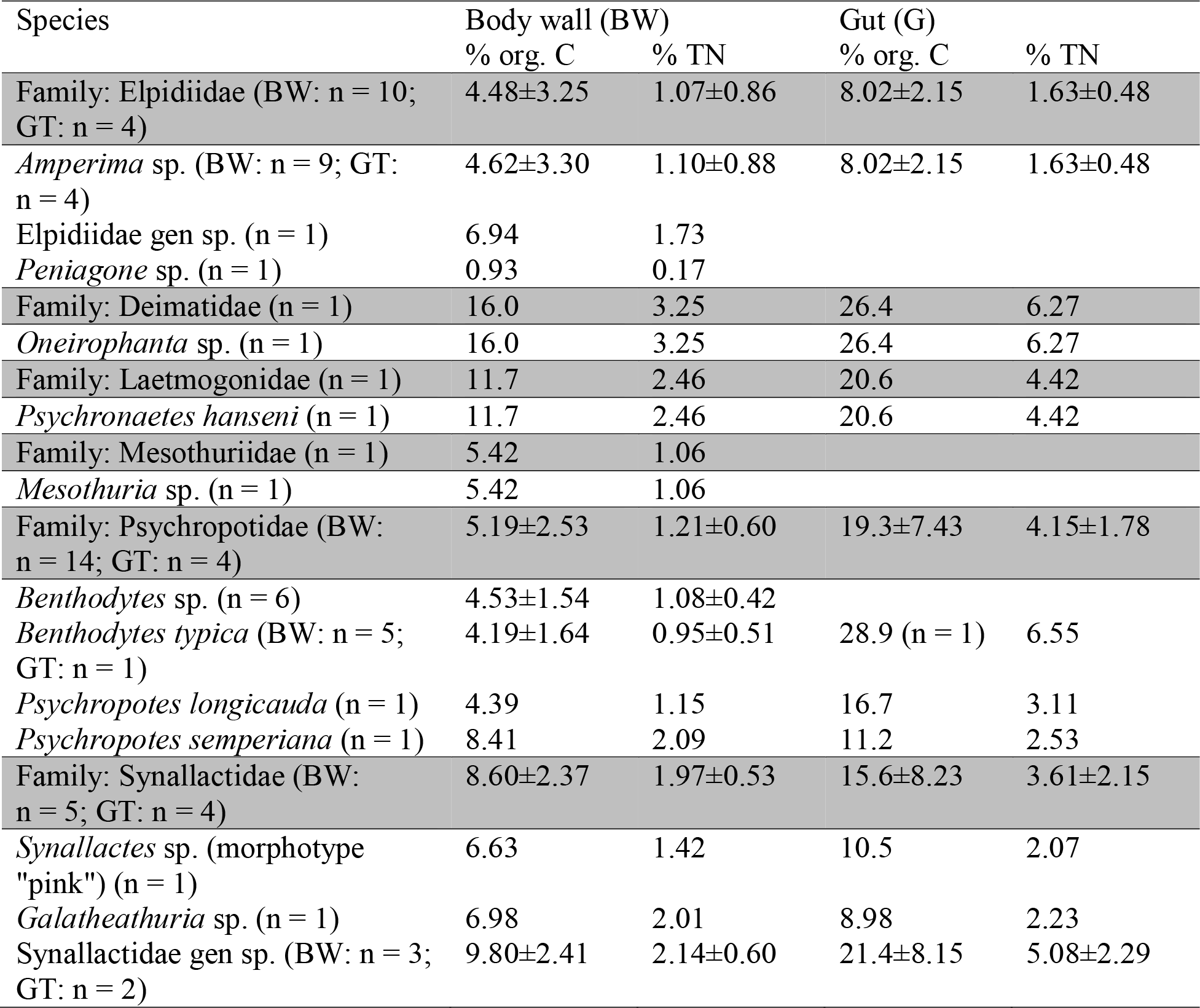
Chemical composition of body wall (BW) and gut tissue (GT) of different holothurian species collected in the Peru Basin. Data are presented as mean±SD.

THAAs, PLFAs, and NLFAs contributed a total 17.4±6.11% to the org. C of all specimens combined (n = 27) and ranged from 1.50 mg C g^-1^ DM THAAs (*Peniagone* sp.; n = 1) to 19.0 mg C g^-1^ DM THAA (*P. hanseni*; n = 1), 0.24 mg C g^-1^ DM PLFAs (*Oneirophanta* sp.; n = 1) to 0.78±0.29 mg C g^-1^ DM PLFAs (*B. typica*; n = 5), and 0.17 mg C g^-1^ DM NLFAs (*Mesothuria* sp.; n = 1) to 2.58±3.32 mg C g^-1^ DM NLFAs (Synallactidae gen sp.; n = 3) (Fig. 3).

**Figure 3.**
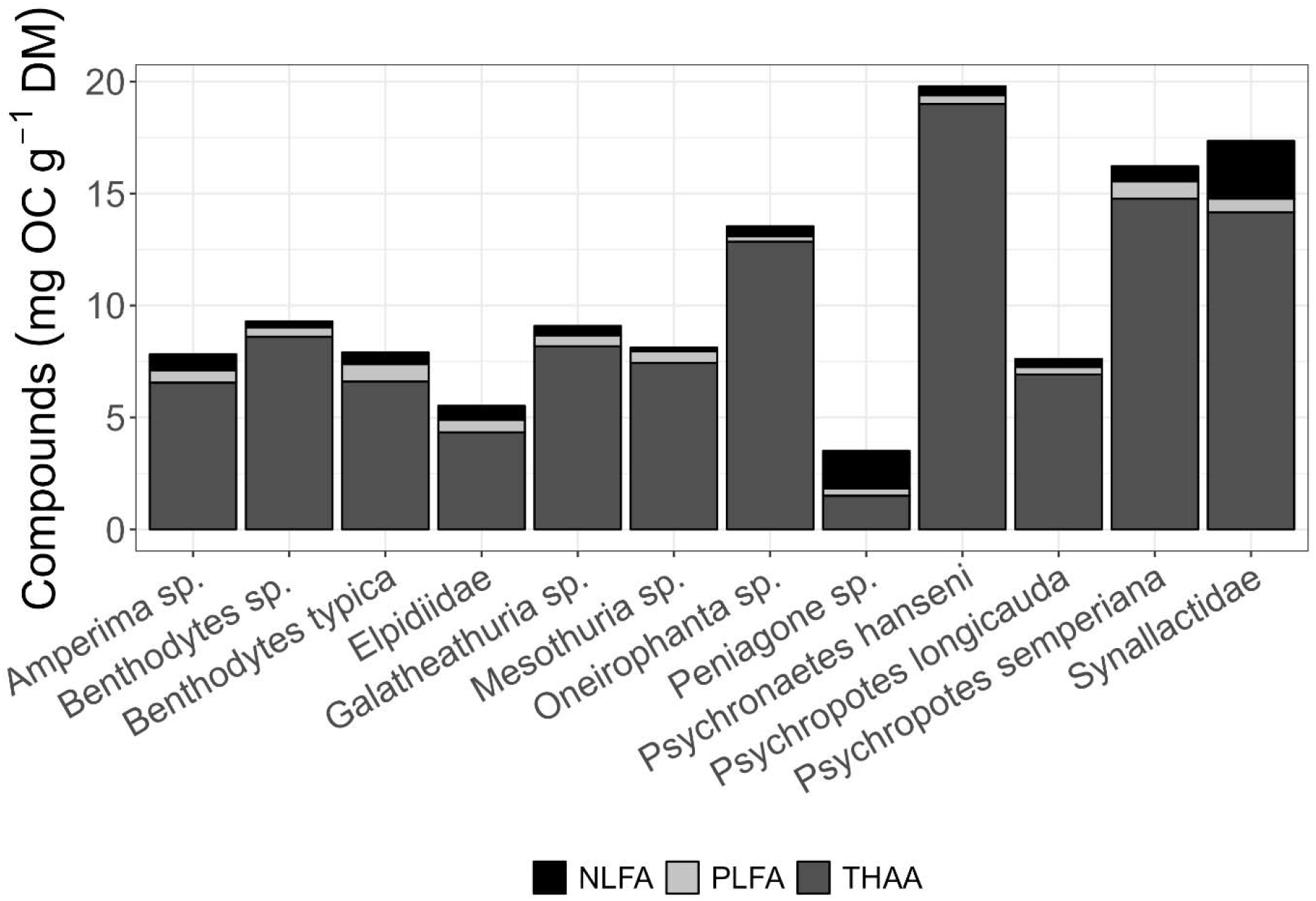
The compounds neutral lipid-derived fatty acids (NLFA), phospholipid-derived fatty acids (PLFA), and total hydrolysable amino acids (THAA) in dried holothurian body walls.

The THAA composition did not differ greatly among species, with the main THAAs in holothurian body walls being alanine, glycine, aspartic acid, and glutamic acid, that contributed between 55.2% (Elpidiidae; n = 1) and 77.4% (*P. longicauda*; n = 1) to total THAAs of body walls (Fig. 4a).

**Figure 4.**
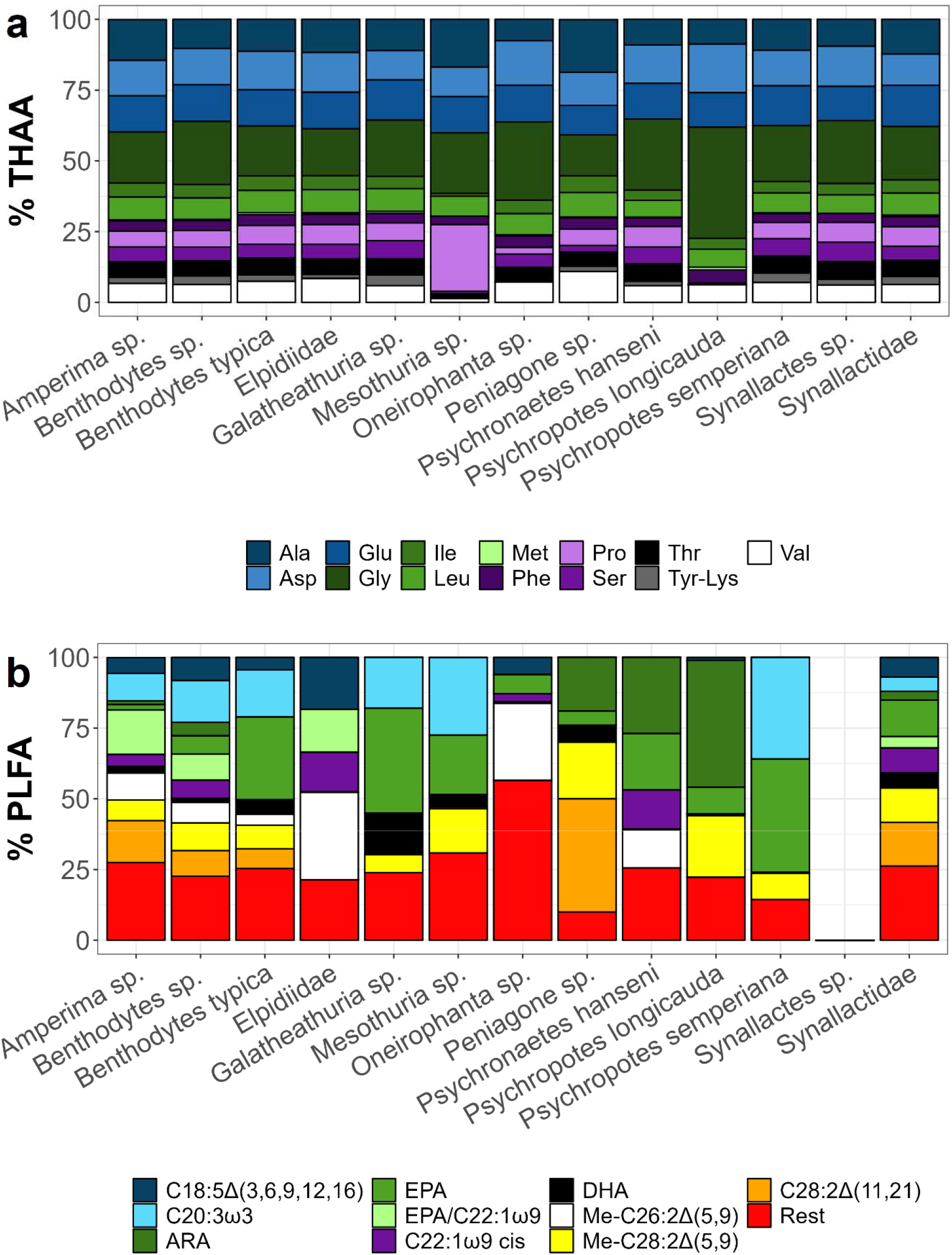
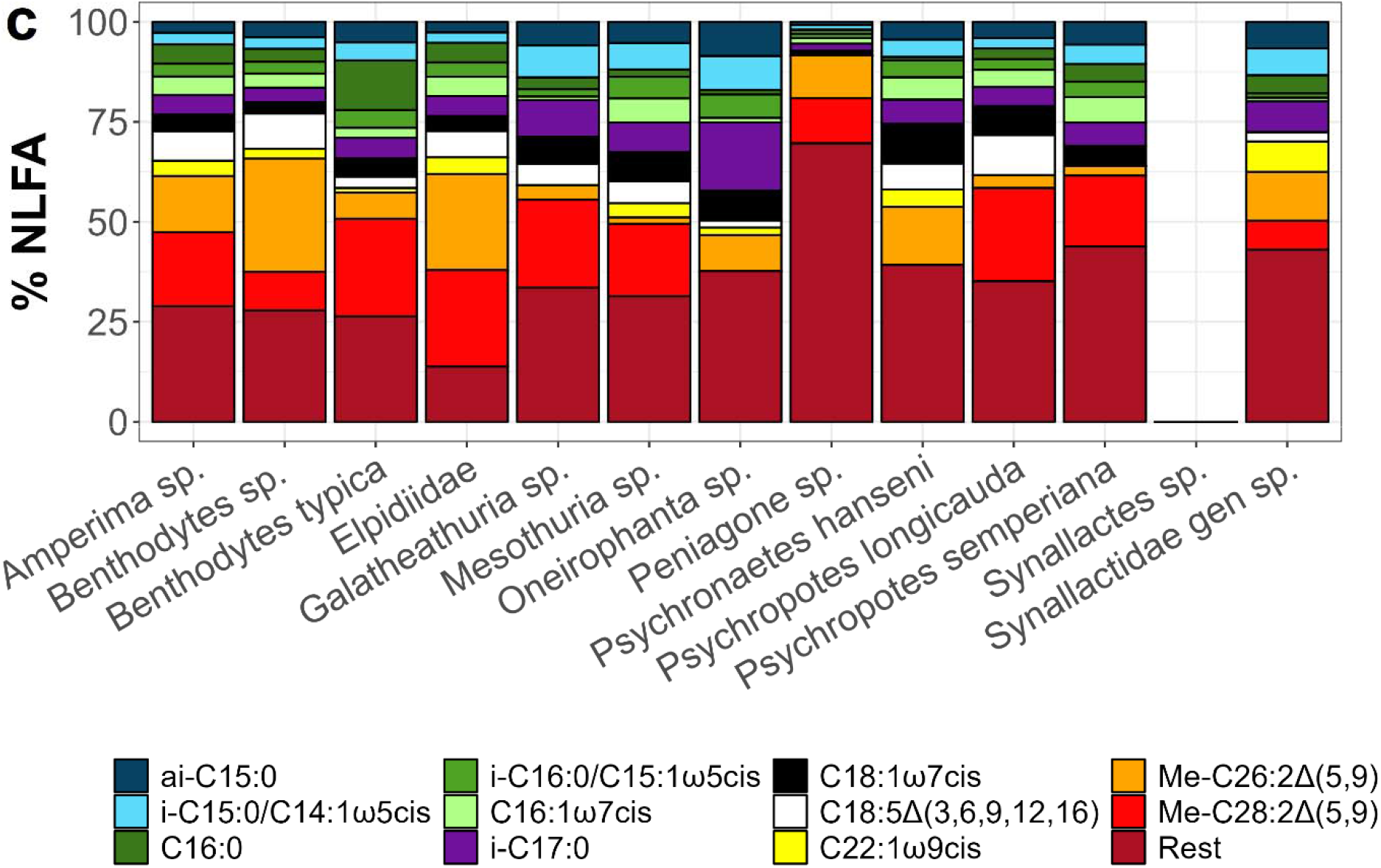
Contribution (%) of individual (a) THAAs, (b) PLFAs, and (c) NLFAs to the total concentrations. The PLFA and NLFA pools ‘Rest’ include all PLFAs and NLFAs, respectively, that contribute <2.5% to total % PLFA and NLFA of the average holothurian tissue.

In contrast, the PLFA (Fig. 4b) and NLFA (Fig. 4c) composition differed strongly between species. Between 10.2% (*Peniagone* sp.; n = 1) and 56.5% (*Oneirophanta* sp.; n = 1) of the PLFAs found in holothurian body walls contributed <2.5% to the total PLFA concentration and were combined as ‘Rest’ (Fig. 4b). The remaining PLFAs consisted to 4.07±7.57% (n = 32) of MUFAs, 10.8±12.6% PUFAs, 32.2±14.1% HUFAs, 9.17±14.0% LCFAs, and 15.5±10.1% methyl-fatty acids. Compared to the average PLFA composition across all holothurian taxa analyzed, Elpidiidae gen sp. (n = 1) had an above average percentage of MUFAs (12.8% of total PLFAs) and methyl-fatty acids (48.1% of total PLFAs). *Oneirophanta* sp. (n = 1) had an above average percentage of SFAs (41.1% of total PLFAs) and *P. longicauda* (n = 1) had an above average percentage of HUFAs (10.0% of total PLFAs).

The NLFAs consisted for 21.6±10.4% (n = 30) of SFA, 10.1±5.80% MUFAs, 5.36±5.03% HUFAs, and 28.5±18.4% methyl-fatty acids (Fig. 4c). Between 13.8% Elpidiidae gen sp. (n = 1) and 69.6% (*Peniagone* sp.; n = 1) of the total NLFAs in holothurian body walls consisted of NLFAs that individually contributed <2.5% to the total NLFA concentration and were therefore combined as ‘Rest’. In comparison to the average NLFA composition across all studied holothurian taxa, *Oneirophanta* sp. (n = 1) had an above average percentage of SFA (41.1% of total NLFAs). *P. hanseni* (n = 1) had an above average percentage of MUFAs (19.9% of total NLFAs), *P. longicauda* had an above average percentage of HUFAs (10.0% of total NLFAs), and Elpidiidae gen sp. had an above average percentage of methyl-fatty acids (48.1%).

The ratio of the essential phospholipid-derived PUFAs EPA to ARA, i.e., the EPA/ARA-ratio, ranged from 0.05±0.13 for *Benthodytes* sp. (n = 6) to 1.75±2.23 for Synallactidae gen sp. (n = 3) (Fig. 2c). In comparison, the ratio of DHA to EPA, i.e., the DHA/EPA-ratio, ranged from 0.01 for *P. hanseni* (n = 1) to 1.27 for *Peniagone* sp. (n = 1) (Fig. 2c). Due to the absence of the PUFAs ARA and/ or EPA in holothurian body wall tissue, no EPA/ARA-ratios were calculated for *B. typica*, Elpidiidae gen sp., *Galatheathuria* sp., *Mesothuria* sp., *Oneirophanta* sp., and *P. semperiana*. Elpidiidae gen sp. lacked both, DHA and EPA, and therefore no DHA/EPA-ratio could be calculated (Fig. 2c).

### Trophic position of holothurians and recycling of amino acids

Holothurians in the Peru Basin had an average δ^13^C-value of -17.4±1.02‰ (n = 31) with a minimum value of -18.4‰ for *Galatheathuria* sp. (n = 1) and a maximum value of -15.5‰ for *Oneirophanta* sp. (n = 1). The average δ^15^N-value was 11.6±1.47‰ (n = 31) with a minimum value of 9.84‰ for *Peniagone* sp. (n = 1) and a maximum value of 14.4‰ for *Oneirophanta* sp. (n = 1) (Fig. 5a).

**Figure 5.**
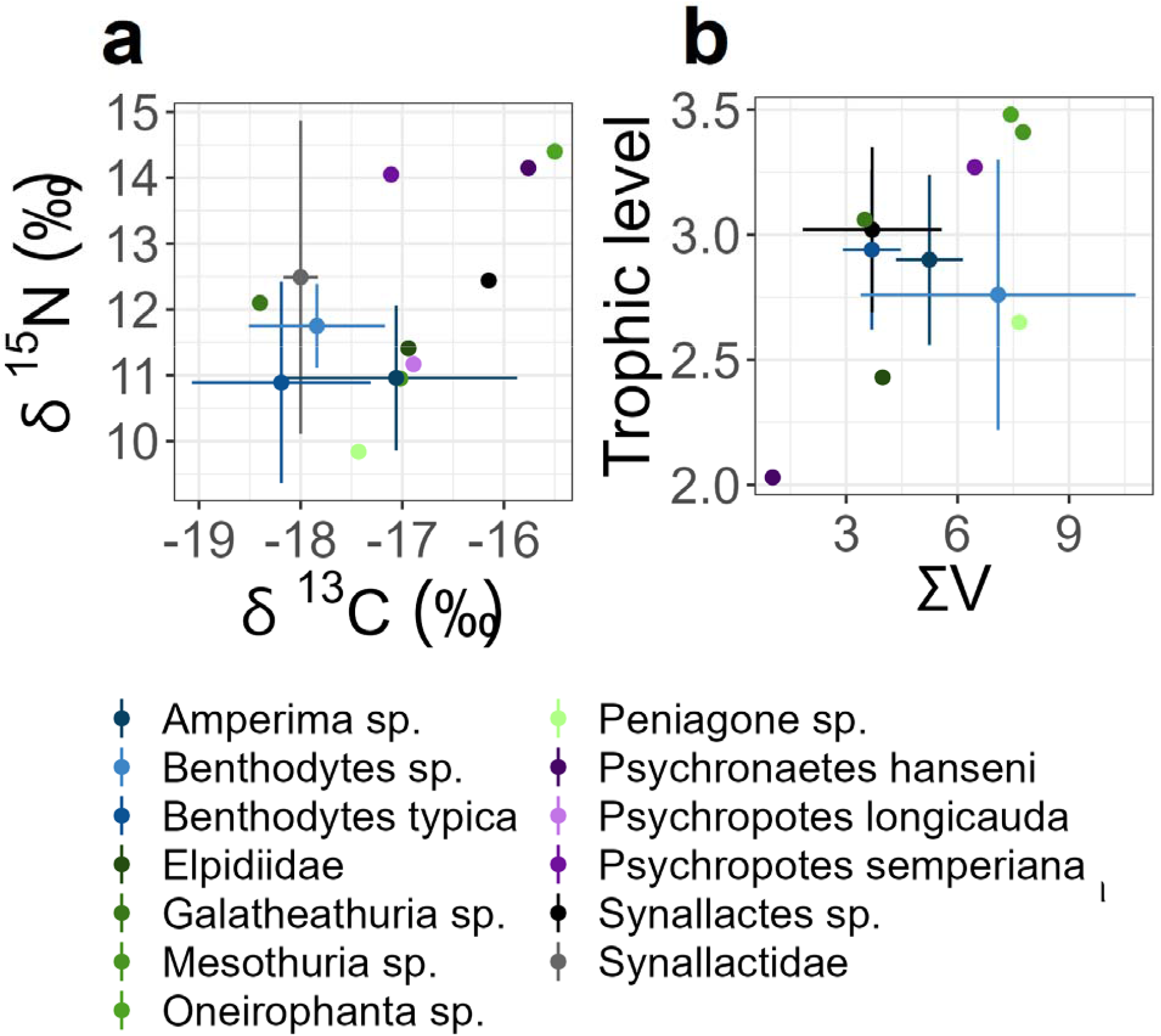
(a) Isotopic composition of carbon (δ^13^C, ‰) and nitrogen (δ^15^N, ‰) of holothurian body wall tissue from the Peru Basin. (b) Trophic position and heterotrophic enrichment factor of holothurians. Error bars in indicate SD.

TL estimates for holothurians in the Peru Basin, based on the δ^15^N values of the THAA glutamic acid and alanine, ranged from 2.0 (*P. hanseni*, n = 1) to 3.5 (*Oneirophanta sp*., n = 1) (Fig. 5b). values ranged from 1.02 to 7.76, with *P. hanseni* having the lowest heterotrophic enrichment and *Mesothuria* sp. having the highest heterotrophic enrichment (Fig. 5b).

## Discussion

### Fatty acid composition of holothurians

Deep-sea megabenthic invertebrates consist to 4.5% DM (cnidarians) to 44.9% DM (crustaceans) of lipids (Drazen et al. 2008a), whereupon holothurians have lipid contents of <1% DM to 5.8% DM (Drazen et al. 2008b). The largest lipid fraction is phospholipids with 14.5% total lipids (crustaceans, Drazen et al. 2008a) to 95.2% total lipids (holothurians; Drazen et al. 2008b). The neutral lipids wax esters and triacylglycerol contribute between <1% (polychaetes) and 83% (crustaceans) to total lipids (Drazen et al. 2008a) and also holothurians consist only of <1% to 2.6% total lipids wax esters and triacylglycerol (Drazen et al. 2008c). Holothurians from the Peru Basin contain between 0.16% DM (*Oneirophanta* sp.) and 2.30% DM (*B. typica*) PLFAs, components of phospholipids, and 0.31% DM (*Oneirophanta* sp.) to 4.71% DM (*Peniagone* sp.) NLFAs, elements of neutral lipids. Hence, they have a relatively high neutral fatty acid content compared to holothurians from Station M (NE Pacific) (Drazen et al. 2008b). This might be related to differences in food availability at the two study sites: The abyssal seafloor at Station M receives on average 22.3 g C m^-2^ yr^-1^ particulate organic carbon (POC) (Baldwin et al. 1998), whereas the POC flux to the Peru Basin is estimated to be 1.49 g C m^-2^ yr^-1^ (Haeckel et al. 2001). As a result, holothurians from the Peru Basin might be adapted to a more food-limited environment by building higher concentrations of storage lipids when they encounter fresh phytodetritus than holothurians at Station M.

### Holothurians trophic level and inferred feeding strategy

Based on the δ^15^N value of body wall tissue, Iken et al., (2001) identified three trophic groups among holothurians from PAP: Group A had δ^15^N values from 10.8 to 12.3‰, group B’s δ^15^N values ranged from 13.2 to 13.9‰, and group C had δ^15^N values from 15.6 to 16.2‰. The δ^15^N values of holothurian tissue from the Peru Basin investigated in this study were lower and ranged from 9.84‰ for *Peniagone* sp. (n = 1) to 14.4‰ for *Oneirophanta* sp. (n = 1). Instead of basing our classification of holothurians from the Peru Basin solely on δ^15^N values, we combined data of trophic level based on compound-specific stable isotope analysis with biomarkers, grain size analysis, and concentration factors for PLFAs.

### Order Elasipodida

*Psychronaetes hanseni* is a deposit feeder of the **family Laetmogonidae**, which has a trophic level of 2.0, low level of heterotrophic re-synthesis of amino acids (∑*V*= 1.02) and feeds selectively (*CF*_*gut content*_ = 25.6) on sedimentary detritus particles of a medium grain size of 17.5 μm which is smaller than the medium grain size of the upper 5 cm of sediment (20.8±0.3 μm; Mevenkamp et al., 2019). Based on the biomarkers present in the body wall tissue of the specimen analysed and in its gut content, parts of the sedimentary detritus likely consists of diatom-derived phytodetritus 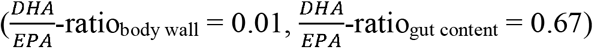.

Elpidiidae gen sp. (**family Elpidiidae**) has a trophic level of 2.4 and medium level of heterotrophic re-synthesis of amino acids (∑*V* = 3.98). This species is a selective deposit feeder (*CF*_*gut content*_ = 14.9) that preferentially feeds upon the PUFA C22:1ω9*cis* which is present in high percentages in its gut content (18%) and in its body tissue (29.0%).

The bentho-pelagic *Peniagone* sp. of the **family Elpidiidae** has an estimated trophic level of 2.7 and a very high level of heterotrophic re-synthesis of amino acids (∑*V* = 7.66). This species has a ‘sweeping’ feeding style (Roberts et al. 2000) and assimilates fresh phytodetritus (Iken et al. 2001) with medium efficiency, as the PLFA concentration in its feces (*CF*_*feces*_) is four times higher than in the surface sediment (this study). In the Peru Basin, *Peniagone* sp. seems to feed on diatom-derived phytodetritus (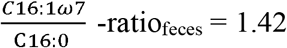; this study).

*Amperima* sp. belongs to the **family Elpidiidae** and its trophic level was estimated to be 2.9±0.3, potentially due to a medium level of heterotrophic re-synthesis of amino acids (∑*V* = 5.24±0.90). This species is a very selective surface deposit feeder (*CF*_*gut content*_ = 58.0) with a ‘sweeping’ feeding style (Roberts et al. 2000) that grazes on very fresh phytodetritus on the surface sediment (Iken et al. 2001). As a result, the gut content of *A. rosea* at PAP has higher concentrations of chlorophyll-*a* compared to surface sediment or phytodetritus (FitzGeorge-Balfour et al. 2010). A more detailed analysis of the phytopigments in this gut content revealed that *A. rosea* at PAP feeds preferentially on cyanobacteria-derived phytodetritus (Wigham et al. 2003). Based on the PLFA composition of its gut content, we found that *Amperima* sp. from the Peru Basin likely feeds on dinoflagellate-derived phytodetritus (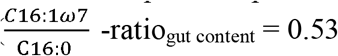= 0.53; this study). Also the body wall fatty acid composition in our study differs substantially from specimens from PAP, as the PLFA profile of PAP specimens is dominated by EPA, DHA, ARA, and C18:0 (Hudson et al. 2004), whereas the PLFA profile of Peru Basin specimens is characterized mostly by EPA co-eluted with C22:1ω9, Me-C26:2Δ(5,9), and C28:2Δ(11,21). Hence, it seems that the feeding niche of the well-studied *Amperima* sp. can differ substantially between ocean basins.

*Benthodytes* sp. from the **family Psychropotidae** has an estimated trophic level of 2.8±0.5 and a very high level of heterotrophic re-synthesis of amino acids (∑*V*= 7.09±3.70). It feeds with a ‘sweeper’ feeding style (Roberts et al. 2000) selectively on smaller sediment particles (medium grain size:13.0 μm) from the surface sediment (medium grain size: 20.8±0.3 μm; Mevenkamp et al., 2019). However, it likely does not or only moderately selects for specifically detritus-enriched particles (*CF*_*gut content*_ = 1.17; *CF*_*feces*_ = 1.81±0.53). In fact, the high percentage of the bacteria-biomarker PLFAs C16:0, C16:1ω7*cis*, and C18:1ω7*cis* in its gut content and feces, and the very high level of heterotrophic re-synthesis of amino acids indicates *Benthodytes* sp. might host a large biomass of living heterotrophic prokaryotes. Unfortunately, in this study no amino acids from gut content or feces were extracted to assess whether this species concentrates detritus that is highly enriched in amino acids. Such an observation was interpreted by Romero-Romero et al., (2021) as a sign that deep-sea holothurians from Station M are secondary consumers of detritus, whereas the microbial community in their guts are primary consumers of detritus. Therefore, we hypothesize that also *Benthodytes* sp. is a secondary consumer, and its microbial gut community is the primary consumer of detritus.

*Benthodytes typica* belongs to the **family Psychropotidae** and its trophic level is estimated to be a bit higher (2.9) than the trophic level of *Benthodytes* sp. This species has a medium level of heterotrophic re-synthesis of amino acids (∑*V*= 3.69) and feeds selectively on smaller particles (medium grain size: 14.2 μm) from the ambient sediment (20.8±0.3 μm; Mevenkamp et al., 2019). These smaller particles contain an at least four times higher concentration of PLFAs than the surrounding sediment (*CF*_*feces*_ = 4.44±1.52) and consist partially of diatom-derived phytodetritus 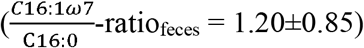. Reliance on phytodetritus is confirmed by the PLFA composition of *B. typica* body walls 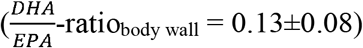. In addition, this species either feeds selectively on sediment-bound prokaryotes or hosts prokaryotes as bacteria-specific PLFAs (i.e., C16:0, C16:1ω7*cis*, and C18:1ω7*cis*) contribute almost 30% to the total PLFA composition in feces, but were not detected in the body wall with >2.5% of total PLFAs. As a medium level of heterotrophic re-synthesis of amino acids was measured, *B. typica* likely has a mixed diet. In this diet, this holothurian species consumes phytodetritus as primary consumer and other types of detritus as secondary consumer following primary processing by a bacterial gut community.

*Psychropotes longicauda* from the **family Psychropotidae** has a medium level of heterotrophic re-synthesis of amino acids (∑*V*= 5.13). Feeding selectively was the highest in our data (*CF*_*gut content*_ = 105), though, surprisingly at PAP this species was found to feed less selectively than *Peniagone diaphana* (FitzGeorge-Balfour et al. 2010). *P. longicauda*’s diet consists likely of diatom-derived phytodetritus 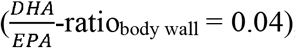, but it is also possible that *P. longicauda* consumes filamentous Rhodophyceae. This algae has been found in gelatinous detritus in the deep sea of the NE Atlantic and Bühring et al., (2002) speculated that *P. longicauda* might feed it sporadically, as the body walls of *P. longicauda* from specimens collected at PAP and in the Peru Basin contain EPA, a PLFA typical for Rhodophyceae, at relatively high concentrations (31% of total PLFA, this study; ∼24% of total fatty acids at PAP (Ginger et al. 2000)).

Additionally, at PAP 70 to 80% of the gut content of this species contained sediment (Iken et al. 2001), which might originate from foraminiferans that Roberts and Moore, (1997) found in its guts together with radiolarians, harpacticoids, nematodes, spicules, and diatoms.

*Psychropotes semperiana* (**family Psychropotidae**) has an estimated trophic level of 3.3, likely related to the high level of heterotrophic re-synthesis of amino acids (∑*V* = 6.46). This species has been classified as surface deposit feeder (Iken et al. 2001) and based on the biomarkers in the body tissue of a specimen collected in the Peru Basin, it consumes diatom-derived phytodetritus (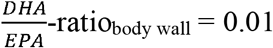, this study).

### Order Holothuriida

*Mesothuria* sp. belongs to the **family Mesothuriidae** and has an estimated trophic level of 3.4. This species could be a subsurface (Iken et al. 2001) or surface deposit feeder (Miller et al. 2000) with a ‘raker’ feeding style (Roberts et al. 2000) or feeding with a ‘wiping’ motion (Hudson et al. 2005). The PLFA composition of its body walls suggests that *Mesothuria* sp. likely consumes diatom-derived phytodetritus 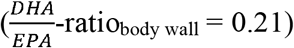. Indeed, in a study on the Hawaiian slope, gut contents of *Mesothuria carnosa* had a 2.7 fold enrichment of chlorophyll a pointing towards selective feeding on phytodetritus (Miller et al. 2000). Furthermore, the very high level of heterotrophic re-synthesis of amino acids (∑*V*= 7.76) from the Peru Basin suggests that *Mesothuria* sp. might also be a secondary consumer of detritus. However, we lack information about its gut content to confirm that it hosts a big(ger) living microbial biomass in its gut that is the primary consumer of detritus.

### Order Synallactida

*Oneirophanta* sp. as member of the **family Deimatidae** has an estimated trophic level of 3.5 and a very high level of heterotrophic re-synthesis of amino acids (∑*V*= 7.43). This species feeds selectively (*CF*_*gut content*_ = 16.6) with a ‘raker’ feeding style (Roberts et al. 2000) and takes up particles with a median grain size of 19.0 μm, which is slightly smaller than the median grain size of sediment particles in the Peru Basin (20.8±0.3 μm; Mevenkamp et al., 2019). The specimen collected in the Peru Basin likely fed on diatom-derived phytodetritus 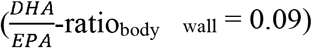 and maybe on bacteria. The very high level of heterotrophic re-synthesis of amino acids and the high trophic level of *Oneirophanta* sp. points to the role of a secondary consumer of detritus, whereas a big biomass of microbial gut community serves as first consumers. However, bacteria-specific PLFAs C16:0, C16:1ω7*cis*, and C18:1ω7*cis*, that were detected in high concentrations in the gut content of *Benthodytes* sp., contribute only 5% to the total PLFA composition in the gut content of *Oneirophanta* sp. Therefore, the diet preferences of this species in the Peru Basin is less clear.

Synallactidae gen sp. (**family Synallactidae**) has an estimated trophic level of 3.0±1.5 and a medium level of heterotrophic re-synthesis of amino acids (∑*V* = 3.70±1.87). It feeds extremely selectively (*CF*_*gut content*_ = 335) and consumes particles of a median grain size (14.5 μm) that is smaller than the median grain size of the surface sediment in the Peru Basin (20.8±0.3 μm; Mevenkamp et al., 2019). The PLFA composition of the body wall and the gut content of Synallactidae gen sp. indicates that this species predates upon agglutinated foraminiferans 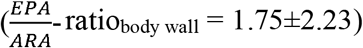 and it consumes diatom-derived detritus (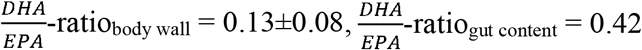, this study). However, it is not possible to differentiate whether Synallactidae gen sp. is a primary consumer of the phytodetritus or a secondary consumer, whereupon the foraminiferans are the primary consumer. The PLFA composition of the feces shows that this holothurian species is also a bacterivore as bacteria-specific PLFAs (i.e., C16:0, C16:1ω7*cis*, and C18:1ω7*cis*) contribute 42% to the total PLFA composition in the feces. If Synallactes hosts a large community of living bacteria, we would expect to detect a significant amount of bacteria-specific PLFAs in the gut content and a higher level of heterotrophic re-synthesis of amino acids. Therefore we assume that Synallactidae gen sp. has a mixed diet consisting of foraminiferans, bacteria, and phytodetritus.

*Galatheathuria* sp. from the **family Synallactidae** has an estimated trophic level of 3.1 and a medium level of heterotrophic re-synthesis of amino acids (∑*V* = 3.50). Similar to Synallactidae gen sp. it feeds extremely selectively (*CF*_*gut content*_ = 285) and *Galatheathuria* sp. seems to consume preferably diatom-derived detritus 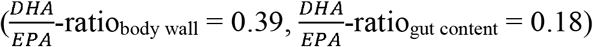.

### Classification of holothurian trophic groups

Here, we propose a classification system of trophic groups for holothurians from the Peru Basin (Fig. 6). It is based on cluster analysis of trophic levels, heterotrophic re-synthesis level of amino acids, feeding selectivity, and diet preferences, instead of on δ^15^N value of body wall tissue. **Trophic group 1** has a trophic level between 2.7 and 3.0, a very diverse diet preference, and includes the species *Amperima* sp., *Benthodytes* sp., *B. typica, Peniagone* sp., and *Psychropotes longicauda*. **Trophic group 2** has a low trophic level of 2.0 to 2.4 and feeds selectively. It includes Elpidiidae gen sp. and *P. hanseni* and **trophic group 3** has a trophic level between 3.0 and 3.5 with a mixed diet and diatom-derived phytodetritus-based diet. It consists of the species *Galatheathuria* sp., *Mesothuria* sp, *Oneirophanta* sp., *P. semperiana*, and Synallactidae gen sp.

**Figure 6.**
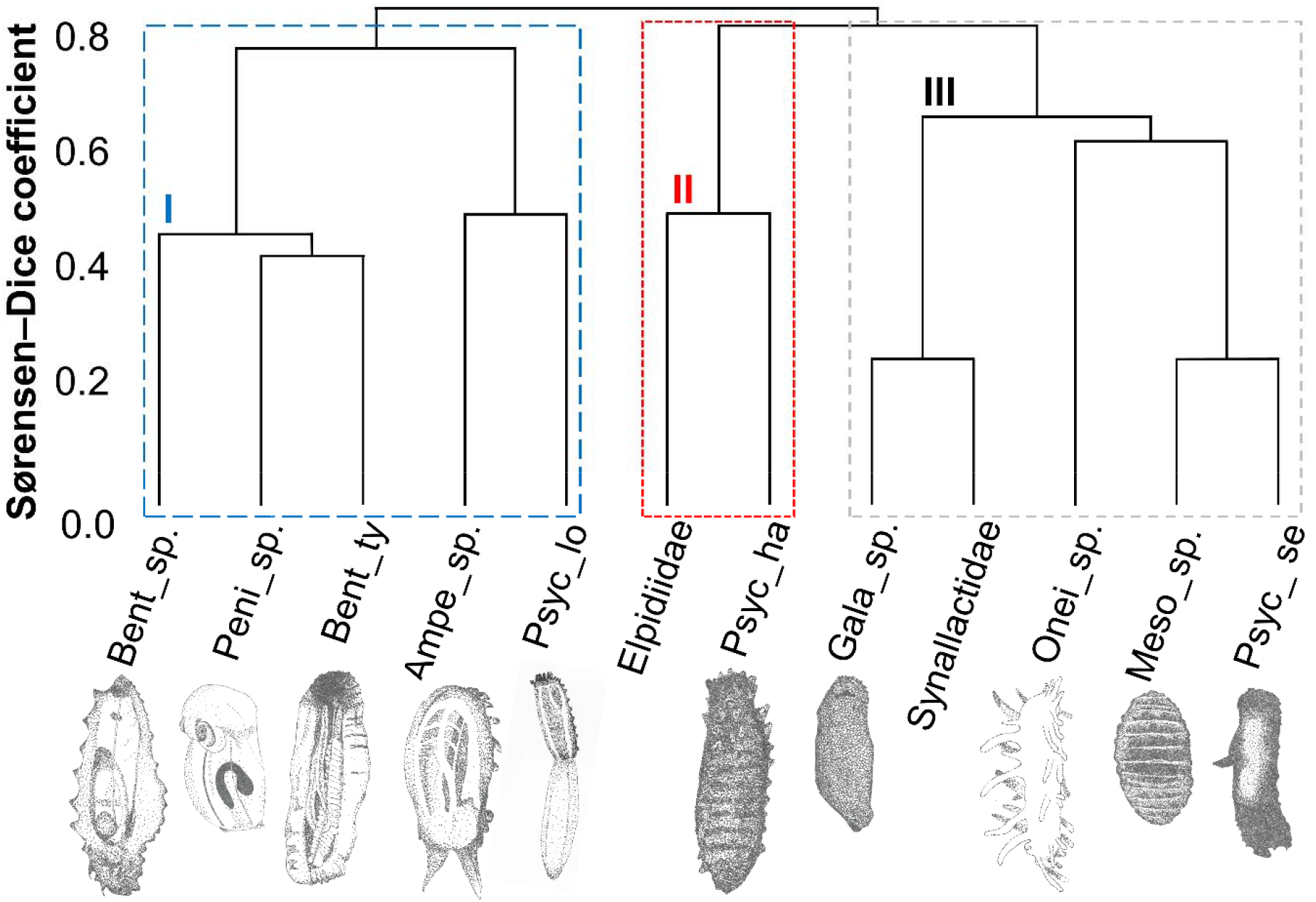
Dendrogram of the Sørensen–Dice coefficient calculated for holothurian species from the Peru Basin based. Trophic group I includes *Amperima* sp. (Ampe_sp.), *Benthodytes* sp. (Bent_sp.), *Benthodytes typica* (Bent_ty), *Peniagone* sp. (Peni_sp.), and *Psychropotes longicauda* (Psyc_lo). Trophic group II comprises Elpidiidae gen sp. (Elpidiidae) and *Psychronaetes hanseni* (Psyc_ha), and trophic group III contains *Galatheathuria* sp. (Gala_sp.), *Mesothuria* sp. (Meso_sp.), *Oneirophanta* sp. (Onei_sp.), *Psychropotes semperiana* (Psyc_se), and Synallactidae gen sp. (Synallactidae). Illustrations of holothurians by Tanja Stratmann.

## Statements and Declarations

### Ethical approval

All applicable international, national, and/or institutional guidelines for animal testing, animal care and use of animals were followed by the authors.

### Sampling and field studies

All necessary permits for sampling and observational field studies have been obtained by the authors from the competent authorities and are mentioned in the acknowledgements, if applicable. The study is compliant with CBD and Nagoya protocols.

### Data availability

The datasets generated during and/or analysed during the current study are available in the PANGAEA repository, **XXX**.

### Author Contribution Statement

TS, DvO conceived the study; TS, DvO performed fieldwork; TS, PvB performed lab analysis; TS drafted the manuscript; TS, DvO, PvB contributed to revising the manuscript to its final version which was approved by all.

## Acknowledgements

The authors thank chief scientist Prof. Antje Boetius, Dr. Felix Janssen, the captain and crew of RV *Sonne*, and the ROV Kiel 6000 team from Geomar (Kiel) for their excellent support during research cruise SO242-2. The authors thank furthermore Dr. Andrey Gebruk for species identification of holothurians. Pieter van Rijswijk, Jana Stratmann, and Jonas Sonntag are thanked for technical assistance during sample processing.

